# Specialized metabolic convergence in the gut microbiomes of cycad-feeding insects tolerant to β-methylamino-L-alanine (BMAA)

**DOI:** 10.1101/2022.12.01.518742

**Authors:** Karina Gutiérrez-García, Melissa R.L. Whitaker, Edder D. Bustos-Díaz, Shayla Salzman, Hilda E. Ramos-Aboites, Zachary L. Reitz, Naomi E. Pierce, Angélica Cibrián-Jaramillo, Francisco Barona-Gómez

## Abstract

Ingestion of the cycad toxins β-methylamino-L-alanine (BMAA) and azoxyglycosides is harmful to diverse organisms. However, some insects are specialized to feed on toxin-rich cycads with apparent immunity. Some cycad-feeding insects possess a common set of gut bacteria, which might play a role in detoxifying cycad toxins. Here, we investigated the composition of gut microbiota from a worldwide sample of cycadivorous insects and characterized the biosynthetic potential of bacteria isolated as putative keystone taxa. Cycadivorous insects shared a core gut microbiome consisting of six bacterial taxa, mainly belonging to the Proteobacteria, which we were able to isolate. To further investigate these potential keystone taxa from diverging lineages, we performed shotgun metagenomic sequencing of co-cultured bacterial sub-communities. We postulate and characterize four putative keystone bacteria from *Serratia, Pantoea*, and two different *Stenotrophomonas* lineages. The biosynthetic potential of these microorganisms includes a suite of biosynthetic gene clusters notably rich in siderophores and carotenoid-like aryl polyene pathways. Siderophore semi-untargeted metabolomics revealed a broad range of chemically related yet diverse iron-chelating metabolites, indicating a complex evolutionary landscape in which siderophores may have converged within the guts of cycadivorous insects. Among these, we provide evidence of the occurrence of an unprecedent desferrioxamine-like biosynthetic pathway that remains to be identified. These results provide a foundation for future investigations into how cycadivorous insects tolerate diets rich in azoxyglycosides, BMAA, and other cycad toxins, and highlight convergent evolution underlying chemical diversity.

## Introduction

Cycads are among the oldest seed plants, with a lineage that traces back to the Permian (Condamine et al., 2015). The genome of one cycad species in the genus *Cycas* was recently published, and with more than 10 Gbp, suggests a series of gene expansions as well as a whole genome duplication prior to divergence from its gingko sister taxa (Liu et al., 2022). These tropical gymnosperms harbor defensive chemicals in their leaves and other tissues, including the azoxyglycoside methylazoxymethanol (MAM), or cycasin, and the non-proteinogenic amino acid β-Methylamino-L-Alanine (BMAA), both of which are toxic to diverse organisms (Liu et al., 2009; Schneider et al., 2002). Previous studies have demonstrated that several insect herbivores are able to detoxify and even sequester the cycad-derived MAM glycoside, cycasin (Nash et al., 1992; Rothschild et al., 1986; Schneider et al., 2002; Teas, 1967), whereas mechanisms of BMAA resistance have not been described. BMAA is capable of blocking glutamate receptors (X. Liu et al., 2009), causing neurotoxic damage in insects (Okle et al., 2013; Zhou et al., 2009) and mammals (Delcourt et al., 2017) as well as developmental problems in *Arabidopsis thaliana* (Brenner et al., 2009). This contrast with the recent observation that BMAA quantities in cycad leaves may not be acutely toxic or deterrent to insects (Whitaker et al., 2022).

In addition to MAM and BMAA, the genomes of a few species of *Cycas* cycads encode for a functional FitD insect toxin that was acquired by horizontal gene transfer from a microbial source, and then expanded into multiple copies (Y. Liu et al., 2022). Many bacteria also have *fitD* genes, as well as the homologous genes *mcf* (‘makes caterpillar floppy’), which have been shown to be broadly distributed and to contribute to insecticidal activity in plant- and nematode-associated bacteria (Ruffner et al., 2015; Flury et al., 2016). These observations, together with BMAA and MAM toxicity, suggest that cycads’ diverse specialized metabolites exert significant evolutionary pressure on cycad-specialized insects and other organisms. Even so, several specialized insect species are able to feed on cycad tissues without adverse effects, including some beetles, thrips, moths, and butterflies (Whitaker & Salzman, 2020), and for at least one group of cycad specialists, *Eumaeus* butterflies, toxin tolerance appears to be a key innovation in the evolution of these insects (Robbins et al., 2021).

The gut bacterial communities of many insects are essential for nutrient acquisition, digestion and detoxification (Jing et al., 2020). Many microorganisms are capable of enzymatically degrading plant specialized metabolites, and growing evidence suggests that symbiotic gut bacteria can help herbivorous insects cope with diets rich in plant defensive chemicals (van den Bosch et al., 2017), and can play a role in resistance to insecticides (Kikuchi et al., 2012; Blanton et al., 2020). An earlier study of the microbiota of cycadivorous insects identified a small core of bacterial taxa that are shared across phylogenetically and geographically distinctive insects (Salzman et al., 2018). The present study aimed to analyze the gut bacteria of a larger sampling of cycad feeding insects, and to explore possible functions by assessing specialized metabolites biosynthesized by key bacterial taxa isolated from the guts of these insects.

To do so, we characterized the bacterial communities of cycadivorous insects from around the world using 16S sequencing, and then applied *EcoMining* (Cibrián-Jaramillo & Barona-Gómez, 2016), a method of capturing bacterial groups and metabolites of interest by co-culturing bacterial sub-communities for shotgun metagenomics, phylogenomics, and integrated omics analyses. The advantage of this method is that it allows for the distillation of functionally relevant fractions of the entire microbiome in response to preconceived biological hypothesis. We then investigated the metabolic functions of potential keystone bacterial taxa by mining for biosynthetic gene clusters (BGCs) predicted to direct the synthesis of specialized metabolites, and confirming the expression of diverse yet related siderophores through semi-untargeted siderophore metabolomics. Our results provide insight into the functional capabilities of key bacteria found in the guts of cycadivorous insects, and their potential importance for chemically-mediated cycad-insect interactions emerging after convergent evolution of iron-chelating metabolites.

## Materials and Methods

### Microbiome global sampling of cycadivorous insects

With the aim of capturing as much ecological and trophic diversity as possible, 12 insect species from 3 orders were collected from the USA, South Africa, China, Australia, Singapore, and Thailand between 2014 and 2017 (**Fig. 1**). Over 90 insect samples were included in the analysis, with detailed collecting information in **Table S1**. All species included in the study are obligate cycad specialists with two exceptions: the neotropical moth, *Seirarctia echo* (Erebidae), which feeds on cycads and plants from multiple families of angiosperms (Whitaker & Salzman, 2020), although the specimens we analyzed were collected from *Zamia* cycads; and the larvae of the African moth *Zerenopsis lepida*, which are obligate cycad folivores in early instars, although some individuals switch to feeding on angiosperms in the fourth instar (Staude et al., 2014). We were able to collect late instar *Z. lepida* feeding on *Encephalartos* and *Stangeria* cycads as well as an angiosperm (*Maesa lanceolata*).

**Fig. 1.**
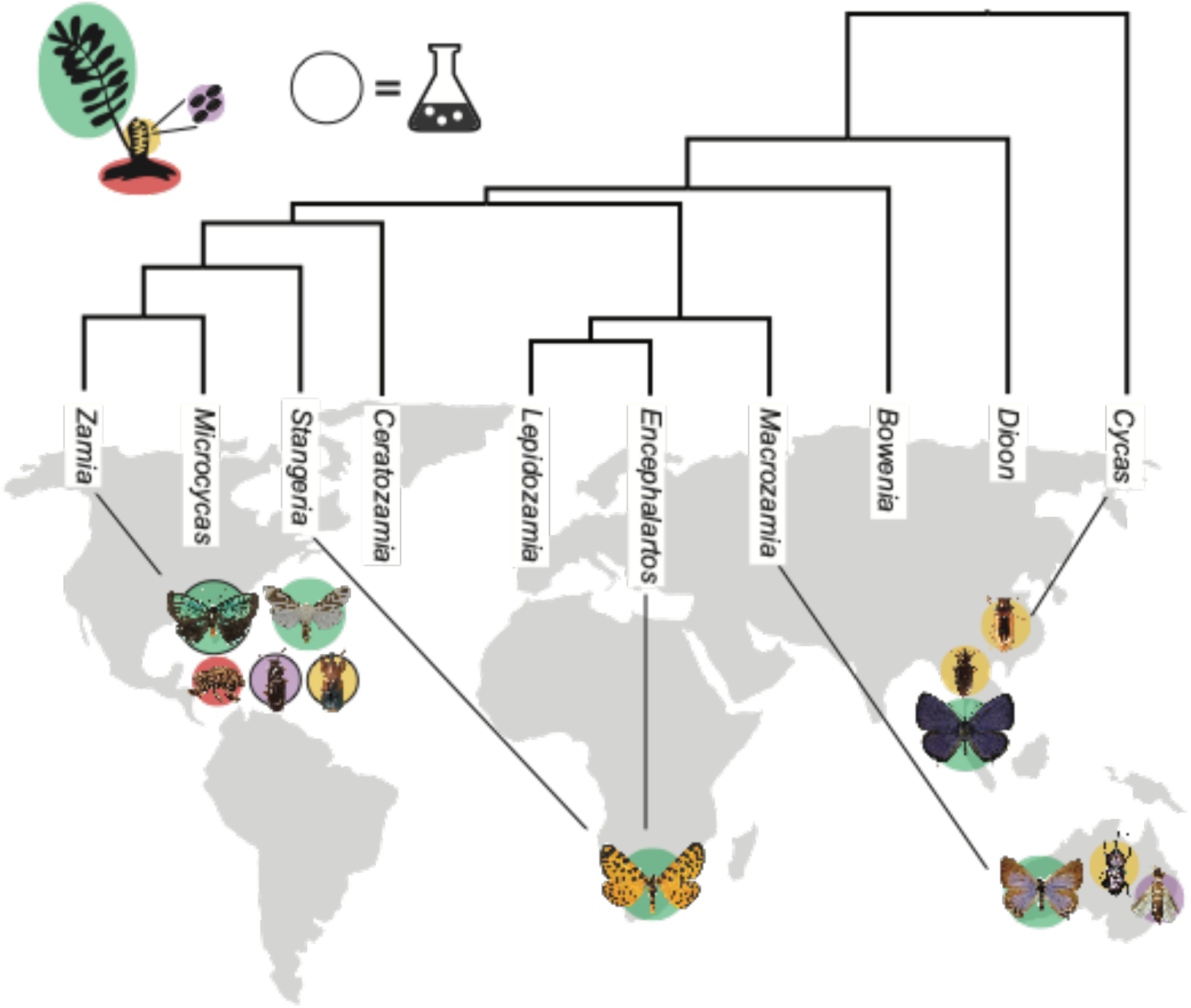
Cycadivorous insect sampling strategy. Coleoptera, Lepidoptera, and Thysanoptera were collected from four continents. Black lines connect each insect species to the cycad genus on which it feeds, and background colors represent the plant tissue on which each insect is specialized: green = leaves, yellow = cones, purple = pollen, and red = roots. All cycad herbivores encounter cycad toxins, regardless of the plant tissue on which they feed. Dark black outlines indicate the three insect genera that were subjected to the *EcoMining* approach.

Insects were processed differently depending on collector, insect type, and size. Lepidopteran larvae in their third instar or older were collected with sterile forceps, submerged in ethanol, and rinsed in sterile PBS. Their guts were then excised using flame-sterilized dissecting tools. Dissected guts were preserved in individual vials containing 97% ethanol. The guts of *Zerenopsis lepida* were not dissected due to their small size; instead, they were surface-sterilized using a 10% bleach solution and preserved whole. Coleopteran specimens were either immediately flash frozen in liquid nitrogen or preserved in ethanol. Whole specimens were then submerged for 5 seconds in a 10% bleach solution and rinsed in PBS for surface sterilization prior to DNA extraction. Finally, specimens from the thysanopteran species *Cycadothrips chadwicki* were neither surface sterilized nor dissected due to their small size; instead, approximately 150 individual thrips were pooled together into a single sample for DNA extraction.

### Amplicon metagenomics of the cycadivorous insect’s gut microbiome

DNA extractions were performed using the Powersoil DNA isolation kit (MoBio Laboratories, Carlsbad, CA, USA) according to the manufacturer’s instructions with the addition of 60 μg of proteinase K to the lysis buffer prior to bead-beating. DNA concentration was assessed using a Qubit Fluorometer (Invitrogen, Carlsbad, CA, USA) and samples with low DNA yields were concentrated as per the DNA extraction kit’s recommendations. DNA extracts were sent to Argonne National Laboratories (Lemont, IL, USA) for library preparation and 16S rRNA gene amplification and sequencing. Amplification of the 16S rRNA was done using barcoded primers 515F (5′-GTGYCAGCMGCCGCGGTAA-3′) and 806R (5′-GGACTACNVGGGTWTCTAAT-3′) of the V4 region (Caporaso et al., 2012), and libraries were pooled and sequenced with Illumina MiSeq instrument in 2×150 paired-end mode. Raw sequences were demultiplexed, filtered, clustered, and given taxonomic assignments following previously published methods (Whitaker et al., 2016). The resulting BIOM table and phylogenetic tree were exported to R and analyzed with the phyloseq (McMurdie & Holmes, 2013), the phyloseq-extended (Mariadassou, 2021), and the microbiome packages. Non-target sequences (mitochondria, chloroplasts, and common lab contaminants) were removed from the dataset and spurious sequences were identified and removed using the PERFect package’s permutation filtering method implemented with the “fast” algorithm (Smirnova, Huzurbazar, & Jafari, 2019). We defined a species’ core microbiome as the set of bacterial operational taxonomic units (OTUs) present in at least 83% of replicates within a species (corresponding to 5 out of 6 replicates, for example). For species represented by multiple life stages (e.g., larva & adult) or, in the case of *Zerenopsis lepida*, multiple host plants, cores were calculated separately for each grouping. For samples with only a single replicate (e.g., the pooled *Cycadothrips* sample), a within-species core could not be obtained, but these samples were included in an across-species core, defined as OTUs present in 98% of all samples.

### Co-culture inoculation for shotgun metagenomics and bacterial isolation

Three insect species were collected and processed to provide inocula for co-cultures: *Rhopalotria slossoni* (Coleoptera), *Pharaxanotha floridana* (Coleoptera), and *Eumaeus atala* (Lepidoptera) (**Table S1**). Twenty-two *R. slossoni* adults, fourteen *P. floridana* adults, and two *E. atala* larvae were surface sterilized by successive washes in 70% ethanol and sterile dd-MilliQ water. The guts of *E. atala* larvae were excised using sterile dissecting tools and then suspended, individually, in sterilized dd-MilliQ water and homogenized with a sterile micropestle. Coleopteran specimens were pooled by species, suspended in sterilized dd-MilliQ water, and homogenized with a sterile micropestle. Resulting biomass from each sample was divided to inoculate two 100 ml co-cultures per sample of liquid BHI medium (Becton Dickinson, Germany) at 0.05x, one without BMAA and one with 20 µM of BMAA, a concentration used in previous studies showing the toxic effects of this metabolite (Berntzon et al., 2013; Popova et al., 2018). Co-cultures were incubated at 23°C with agitation at 150 r.p.m for 48 hrs, after which each co-culture was used to inoculate petri dishes containing different bacterial media described below. DNA was extracted from the eight co-cultures for shotgun metagenomics. Two additional co-cultures (BMAA +/-) were generated from the pooled *P. floridana* biomass and were used for genomic isolation but not shotgun metagenomics.

### Shotgun metagenome sequencing and taxonomic analysis

DNA was extracted from co-cultures using a standard phenol-chloroform protocol (Sambrook & Russell, 2006), and DNA quality was checked by Nanodrop 2000/2000c (Thermo Scientific, USA). Libraries were prepared with the Truseq nano kit and sequenced at UGA, Cinvestav (Irapuato, Mexico) using the NextSeq Illumina platform 2×150 paired-end reads format. Obtained reads were checked with FastQC v0.11.9 (Wingett & Andrews, 2018) and low-quality bases were trimmed with Trimmomatic v0.32 (Bolger et al., 2014). Metagenomic profiling of high-quality paired reads was performed using kraken2 (Wood et al., 2019) with the maxikraken database (September 2018 release; available at https://lomanlab.github.io/mockcommunity/mc_databases.html). Abundance and taxonomic lineage tables were obtained from the standard kraken2 output by using an in-house script, then exported to R to be analyzed with the phyloseq (McMurdie & Holmes, 2013), the taxize (Chamberlain & Szöcs, 2013), and the microbiome packages.

### Shotgun metagenome assembly, mining, and binning

Assembly of the six metagenomes obtained was performed with metaSPAdes v3.10 using default parameters (Nurk et al., 2017). The resulting scaffolds were annotated using RAST (Aziz et al., 2008), taxonomically classified using kraken2 with the same database and parameters used for read classification, and mined using antiSMASH v5.0 (Weber et al., 2015), BiG-SCAPE and CORASON (Navarro-Muñoz et al., 2020). To obtain the core biosynthetic gene clusters (BGCs), each complete and non-redundant BGC obtained from antiSMASH was used to construct a presence/absence matrix. A core BGCs plot was then created using the UpSetR R package v1.4.0 (Lex et al., 2014), with all the set intersections in this matrix. Scaffolds were binned with the PATRIC Metagenomic Binning Service (Parrello et al., 2019) using default parameters, from which nine metagenome-assembled genomes (MAGs) were obtained: 4 for *Stenotrophomonas*, 4 for *Serratia*, and 1 for *Pantoea* (**Table S2)**. To obtain 16S sequences from the metagenomes we used the 16S sequence recovery pipeline of Anvi’o (Eren et al., 2015).

### Isolation and identification of bacteria from co-cultures in semi-selective media

Semi-selective media were chosen to target specific bacterial phyla that are known to be represented in cycadivorous insects’ microbiomes (Salzman et al., 2018). These were: 1) SFM media, (mannitol: 20 g/L; soya flour 20 g/L; and agar 17 g/L for solid media); 2) ISP4 and ISP4N-, (starch: 10.0 g/L; dipotassium phosphate: 1 g/L; magnesium sulfate: 1 g/L; sodium chloride: 1 g/L; ammonium sulfate: 2 g/L for ISP4 media, none for ISP4N-media; calcium carbonate: 2 g/L; ferrous sulfate: 1 mg/L; magnesium chloride: 1 mg/L; zinc sulfate: 1 mg/L; final pH 7.0; and agar 17 g/L for solid media); 3) BHI media at concentrations 0.02x and 0.05x (Becton Dickinson, Germany); 4) FLA media (Fortified lipid agar) (tryptic soy broth: 16 g/L; vegetable oil: 10 mL/L; nutrient broth 12 g/L; yeast extract: 5 g/L; and agar 17 g/L for solid media); and 5) NTBA media (peptone: 5 g/L; beef extract: 3 g/L; bromothymol blue: 0.025 g/L; 2,3,5-triphenyl tetrazolium chloride: 0.04 g/L; and agar 17 g/L for solid media).

The Petri dishes were incubated at 22°C for 72 hrs. After this, only unique bacterial morphotypes were selected for further isolation and purification in their corresponding media. Once bacterial morphology was homogeneous on each isolation plate, isolates were grown on 100 ml of their corresponding media for 48 hrs. DNA extraction and quality measurements was done as before. gDNA was used for PCR amplification of the 16S rRNA region using the universal bacterial primers 27F and 1492R (LANE & D. J, 1991). The amplification yielded PCR fragments of 1.4 Kbp in length, which were purified with the Qiagen QIAquick PCR purification kit (Hilden, Germany) and sequenced using Sanger chemistry at UGA, Cinvestav (Irapuato, Mexico). Taxonomic identity of the sequences was determined using Blastn against the SILVA rRNA database v132 (Quast et al., 2013). Each sequence was classified according to the best hit (>97% sequence identity) (**Table S3**). An alignment of all of the obtained 16S rRNA sequences was constructed with ClustalW (Thompson et al., 2002) and edited with Gblocks v0.91 to remove ambiguous positions (Castresana, 2000). A phylogenetic tree was constructed using MrBayes v3.2 (Ronquist & Huelsenbeck, 2003), with a 4by4 nucleotide substitution model for 100 000 generations, with sampling every 100 generations. The graphical representation of the phylogenetic trees was obtained with Iroki Tree Viewer (Moore et al. 2020).

### Genomic sequencing and natural products mining of isolated strains

We selected six strains of interest from the isolates based on two criteria: 1) their prevalence in the gut microbiomes of *R. slossoni, P. floridana*, and *E. atala*; and 2) their biosynthetic potential to produce conserved natural products. For genomic characterization of these strains, we grew monocultures of the selected *Serratia, Pantoea*, and *Stenotrophomonas* strains, and the obtained biomass was used for DNA extraction and quality measurements 0as before. Libraries were prepared using the Truseq nano kit and sequenced at UGA, Cinvestav (Irapuato, Mexico) using the NextSeq Illumina platform with 2×150 paired-end reads format. Raw reads were checked with FastQC v0.11.9 and low-quality bases were trimmed using Trimmomatic v0.32 (Bolger et al., 2014). Multiple *de novo* genomes were assembled with SPAdes v3.15.3 using default parameters (Zerbino & Birney, 2008) and annotated with RAST (Aziz et al., 2008) (**Table S2**). CORASON (Navarro-Muñoz et al., 2020) was used to mine turnerbactin-like siderophores in the different MAGs and genomes within a phylogenetic context. CORASON hits that contained homologs of the turnerbactin-like non-ribosomal peptide synthetase (NRPS) were obtained using a BlastP (e-value and cut-off of 0.001).

### Siderophore genome mining

To further characterize the putative siderophore BGCs detected by antiSMASH v5.0 (Weber et al., 2015), we functionally annotated genes in the BGCs that were previously implicated in iron metabolism. This included enzymes annotated as acyl CoA-dependent acyltransferases, FAD-dependent amine monooxygenases and iron reductases, as well as TonB-dependent receptors and ABC transporters. The presence of conserved DNA motifs corresponding to iron boxes, known to be involved in iron regulation in bacteria (Hantke, 2001), were also determined and used to mine the genomes. An iron box database was constructed using DNA motifs previously described for Gram-positive bacteria (TTAGGTTAGGCTCACCTAA and TGATAATNATTATCA) (Baichoo & Helmann, 2002; Cruz-Morales et al., 2017), and Gram-negative bacteria (GATAATGATAATCATTATC, GATAAAATTAATCAGCCTC, and ATTAATAAAAACCATTGTC, GATAATGAGAATCATTATT, GATAATTGTTATCGTTTGC, TATAATGATACGCATTATC, TGTAATGATAACCATTCTC, GAATATGATTATCATTTTC, GAAAATGATAATCATATTC, ATAAATGATAATCATTATT, GATAATCATTTTCAATATC) (Baichoo & Helmann, 2002; Sauvage et al., 1996; Panina et al., 2001). The presence of iron box motifs was analyzed with PREDetector v3.1 (Tocquin et al., 2016) using the iron box motifs dataset.

### Phylogenomic analysis of putative keystone taxa

To establish the relationships between MAGs and genome sequences of bacterial isolates, phylogenies of *Stenotrophomonas, Pantoea*, and *Serratia* were constructed using representative genomes from each genus, along with our newly generated genome sequences and recovered MAGs (**Table S4, S5, and S6**). For each phylogeny, a protein core was obtained using Get-Phylomarkers (Vinuesa et al., 2018), computed with Get-Homologues using the algorithms bidirectional best-hit (BDBH), Clusters of Orthologous Groups-triangles (COGtriangles), and OrthoMCL (Markov Clustering of orthologs, OMCL) with default parameters (**Table S7, S8, and S9**). Resulting matrices, containing 39, 64, and 712 proteins, respectively, were used to construct phylogenetic trees with MrBayes v3.2 (Ronquist & Huelsenbeck, 2003) with a mixed substitution model based on posterior probabilities (aamodel[Wag]1.000) for proteins for 100 000 generations. The trees were visualized with FigTree v1.4.2 (Rambaut, 2014).

### Identification of siderophores produced by keystone bacteria through HPLC and MS

The *Pantoea, Serratia*, and *Stenotrophomonas* bacteria were subjected to semi-untargeted siderophore extraction and characterization. Each strain was cultivated in two separate Erlenmeyer flasks containing LB medium, with and without 200 µM of the iron-chelator compound 2,2′-dipyridyl (DIPY) for four days, as previously (Cruz-Morales et al., 2017). Culture growth was checked by OD_600_ measurements every 12 hrs and cultures were harvested upon reaching saturation. Cultures were then centrifuged at 8 000 rpm and the supernatant was collected and lyophilized. Dry extracts were resuspended in 10 ml of sterile dd-MilliQ water and separated in two Falcon tubes, each with 5 ml of the resuspended extracts. Iron-chelating metabolites were converted to their ferric complexes by addition of 1 M FeCl_3_ to one of the tubes of each condition. All samples were then concentrated by filtering the supernatants with Millipore Millex-GN filters with a nylon membrane of 0.22 µm. Samples were analyzed on a Thermo Ultimate 3000-uHPLC instrument equipped with a quaternary pump, a diode array detector and a ZORBAX Eclipse XDB-C18 (Agilent, USA) analytical column (4.6×150mm, 5um). Analytical conditions were as follows: The mobile phase comprised a binary system of eluent A, 0.1% trifluoroacetic acid, and eluent B, 100% acetonitrile. The run consisted of a gradient from 0 to 100% B for 25 min. Tris-hydroxamate-Fe^3+^ complexes, formed after the addition of iron chloride to bacterial extracts, were detected at a wavelength of 435 nm. Fractions were collected and pooled by bacterial genus for mass spectrometry (MS) analysis in an ion trap LTQ Velos (Thermo Scientific, Waltham, USA). MS/MS analysis of selected ions was performed with a collision energy of 20 eV, and spectra were analyzed using the XCalibur software v3.0 (Thermo Scientific, USA). Metal-chelating metabolites were identified by comparing fragmentation patterns with those of siderophores previously reported (Cruz-Morales et al., 2017) or included on the Bertrand’s siderophore database (http://bertrandsamuel.free.fr/siderophore_base/index.php) and the GNPS database used to generate GNPS feature-based molecular networks (Wang et al., 2016; Aron et al., 2020).

## Results

### 16S taxonomic profiling of cycadivorous insects’ guts identifies a global microbiome core

16S amplicon sequencing of DNA extracted from the guts of a global sampling of cycadivorous insect species yielded 6 317 577 bacterial reads. After removing non-target and spurious OTUs, we were left with 5 027 153 reads across 92 samples. Observed bacterial species richness ranged from 41 to >700 OTUs within a single insect sample. A total of 26 bacterial phyla were detected (**Fig. 2A**), with Proteobacteria comprising moderate to dominant portions of the microbiomes of most insects with a few exceptions: *Zerenopsis lepida* larvae feeding on *Stangeria* cycads (but not other plants) were dominated by Firmicutes, *Tychiodes* sp. adults were dominated by Tenericutes, and *Seirarctia echo* larval microbiomes were composed of relatively equal proportions of Proteobacteria, Bacteroidetes, and Firmicutes. Ordinations based on unweighted UniFrac distances of the bacterial communities reveal clustering according to insect species, life stage, and host plant, though with some overlap in overall community composition across insects/groupings (**Fig. 2B**). Insects’ core microbiomes ranged widely in richness, from 17 core OTUs in *Luthrodes pandava* larvae (n = 7) to 185 OTUs in *Eubulus* sp. larvae (n = 7). In general, the core microbiomes of Coleoptera tended to be larger than in Lepidoptera. A cross-species core included 6 OTUs found in all insect species. These OTUs belong to the genera *Acinetobacter* (OTU 3), *Pantoea* (OTU 7), *Enterobacter* (OTU 2), *Enterococcus* (OTU 4), *Lactobacillus* (OTU 154), and *Stenotrophomonas* (OTU 6). These taxa were then placed within a phylogenetic context after isolation and analysis of keystone bacterial species as described in the following sections (**Fig.3**).

**Fig. 2.**
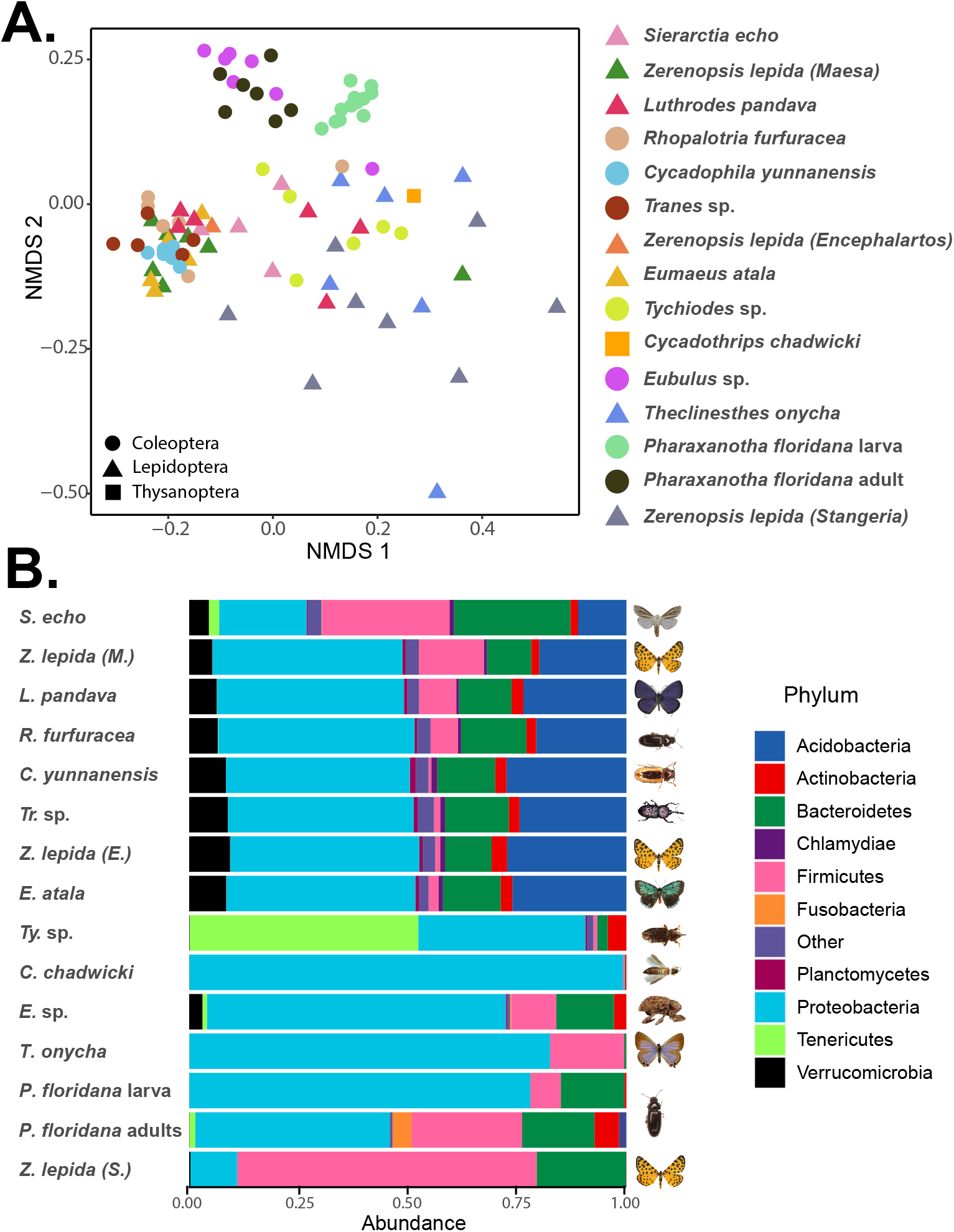
Composition and distribution of cycadivorous insect’s gut microbiome. **A**. Ordination of community composition among a global sampling of cycad-feeding insects. In an NMDS plot of unweighted UniFrac distances, samples cluster to varying degrees by insect species, life stage, and host use. Sample libraries were rarefied to 5 000 sequences (keeping samples with >1 000 sequences) prior to ordination. **B**. Proteobacteria dominate the gut microbiomes of most cycadivorous insects, followed by Firmicutes and Bacteriodetes.

**Fig. 3.**
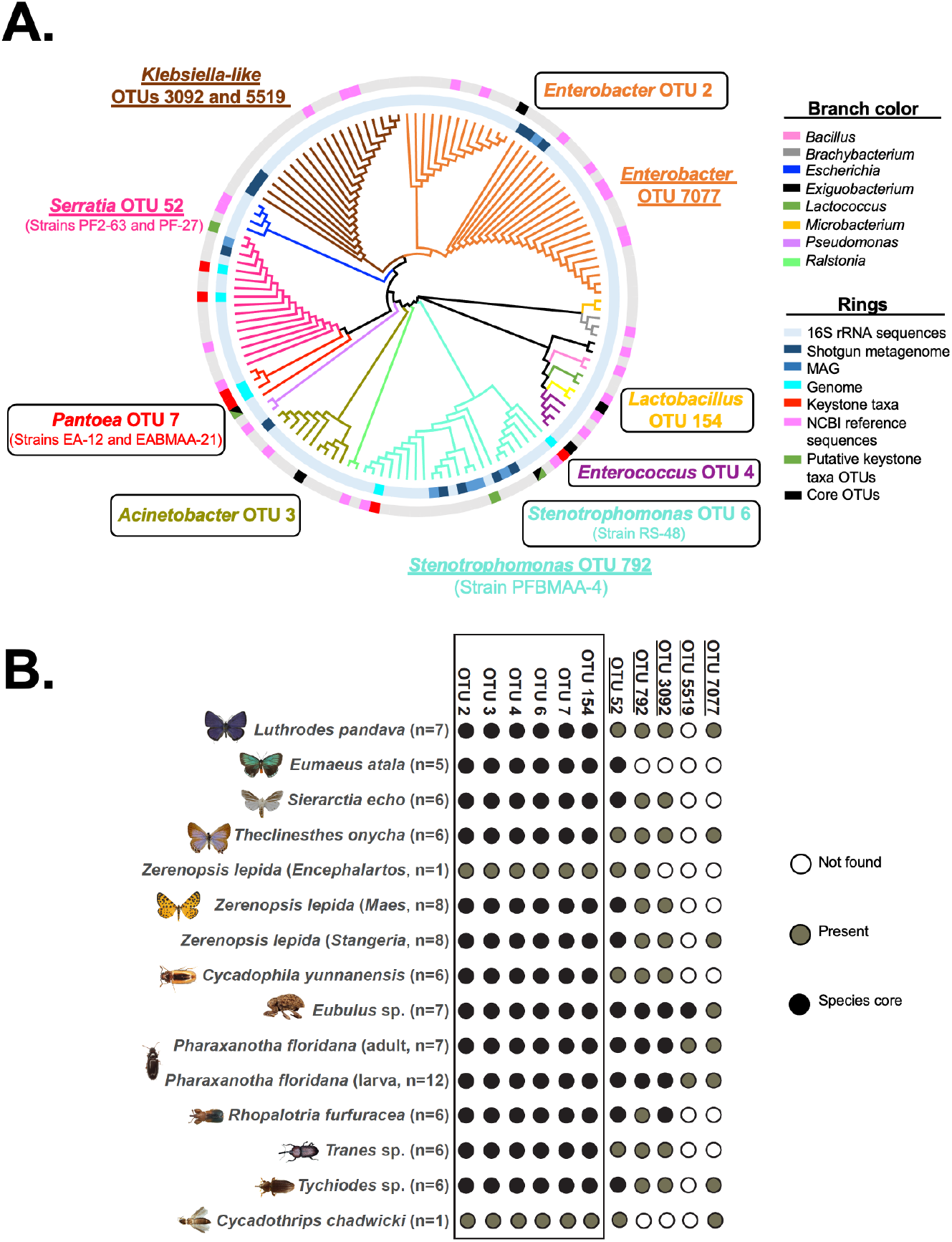
Identification and isolation of proposed keystone strains for functional analysis. **A**. 16S rRNA phylogenetic tree including 79 bacterial strains isolated from the guts of *Eumaeus atala, Pharaxanotha floridana*, and *Rhopalotria slossoni*. The 16S rRNA gene from the six genomes of the genera *Serratia, Pantoea*, and *Stenotrophomas*, plus all the metagenomes data and the 16S amplicon data were included. Proposed keystone taxa identified after the integration of different datasets (panel B) are highlighted: core taxa are shown within boxes and semi conserved taxa shown underlined. Specific strains selected for further experimental characterization are shown between parentheses below their cognate OTUs. **B**. Metagenomics and genomics data integration reveal keystone taxa. Black circles indicate OTUs belonging to an insect species’ core, grey circles represent non-core OTUs present in some individuals within a species/grouping, and white circles indicate OTUs that were not found in any individual within a species/grouping. As in panel A, core taxa are shown within a box and semi conserved taxa shown underlined without a box.

### Isolation of putative keystone taxa from the core microbiome of cycadivorous insects’ guts

To direct bacterial isolation towards ecologically relevant strains from the broad diversity of cycadivorous insects investigated, we adopted the *EcoMining* approach, which we have successfully employed to investigate the microbiome of cycad’s coralloid roots and its biosynthetic potential (Gutiérrez-García et al., 2019). We obtained bacterial isolates from the sub-community co-cultures of the dissected guts of *Eumaeus atala, Pharaxanotha floridana*, and *Rhopalotria slossoni* (**Fig. 1**, circles). This approach led to two types of cultures for further analyses: (i) bacterial isolates obtained in selective and semi-selective media, which were characterized after amplification and sequencing of their 16S rRNA gene, and eventually their whole genome for selected strains; and (ii) sub-community co-cultures grown under functionally relevant conditions, such as the presence of the cycad toxin BMAA, which were shotgun sequenced to identify key taxonomic and functional genes, both at the single-gene level and as part of metagenome-assembled genomes (MAGs) (**Fig. 3A, Table 1**).

**Table 1.** Metagenomes and genomes generated in this study.

| (Meta)genome sequence / strain | Type | Bacterial genus | Origin<sup>a</sup> | Accession number<sup>b</sup> |
| --- | --- | --- | --- | --- |
| EA | Metagenome | NA | EA |  |
| EABMAA | Metagenome | NA | EA + BMAA |  |
| PF | Metagenome | NA | PF |  |
| PFBMAA | Metagenome | NA | PF + BMAA |  |
| RS | Metagenome | NA | RS |  |
| RSBMAA | Metagenome | NA | RS + BMAA |  |
| Pa-EAmG | MAG | *Pantoea* | EA |  |
| Se-PFmG | MAG | *Serratia* | PF |  |
| St-PFmG | MAG | *Stenotrophomonas* | PFBMAA |  |
| Se-PFBMAAmG | MAG | *Serratia* | PFBMAA |  |
| St-PFBMAAmG | MAG | *Stenotrophomonas* | PFBMAA |  |
| Se-RSmG | MAG | *Serratia* | RS |  |
| St-RSmG | MAG | *Stenotrophomonas* | RS |  |
| Se-RSBMAAmG | MAG | *Serratia* | RSBMAA |  |
| St-RSBMAAmG | MAG | *Stenotrophomonas* | RSBMAA |  |
| EA-12 | Genome | *Pantoea* | EA |  |
| EABMAA-21 | Genome | *Pantoea* | EABMAA |  |
| PFBMAA-4 | Genome | *Stenotrophomonas* | PFBMAA |  |
| RS-48 | Genome | *Stenotrophomonas* | RS |  |
| PF2-63 | Genome | *Serratia* | PF2 |  |
| PF-27 | Genome | *Serratia* | PF |  |

NA, not applicable; EA, *Eumaeus atala*; PF, *Pharaxanotha floridana*; RS, *Rhopalotria slosoni*. <sup>a</sup> The origin refers to the co-culture from which the metagenome DNA was extracted and sequenced or the bacterial strain isolated and subsequently sequenced. <sup>b</sup> Submission in progress.

To match the different datasets generated, we extracted a subset of the 16S profiling data from *Rhopalotria furfuracea* adults (n = 6), *Pharaxanotha floridana* adults (n = 7), and *Eumaeus atala* larvae (n = 5), for comparison to the taxonomic information generated by co-culture shotgun metagenomic sequencing. Alpha diversity analysis showed that shotgun metagenomes recovered more OTUs than 16S amplicon sequencing (**Fig. S1A**) and that these OTUs belonged to fewer phyla, with Proteobacteria being the most dominant phylum, reflecting a partial but marginal growth bias of the co-cultures (**Fig. S1B**). These analyses also showed that addition of BMAA to the co-cultures does not alter the taxonomic composition of bacterial communities; instead, host species identity was the biggest factor contributing to taxonomic dissimilarities among sub-communities (**Fig. S1A**). The fact that BMAA did not alter the microbial diversity of the co-cultures is congruent with the fact that the isolated sub-communities are adapted to this toxin present in the insect’s diet.

The Proteobacteria diversity recovered from the shotgun metagenomes and their corresponding 16S sequences revealed the presence of the genera *Serratia, Acinetobacter, Stenotrophomonas*, and *Enterobacter* across all three insect species regardless of the metagenomic approach, albeit with different patterns of prevalence (**Fig. 3B)**. Given that Proteobacteria was also the most abundant phylum in the 16S amplicon dataset, it is reasonable to conclude that this is a constitutive part of the insects’ gut microbiomes. We also detected some uncultured bacteria from the Enterobacteriaceae family, present in the metagenomes obtained from *Pharaxanotha floridana*, with and without BMAA (PF and PFBMAA, respectively), and *Rhopalotria slossoni* (RS). Based on the 16S phylogeny, the closest match to these previously uncultured bacteria belongs to the genus *Klebsiella* (OTUs 3092 and 5519), with sequence identity scores between 90% and 93% depending on the metagenome.

Using this integrated dataset as a guide, we then selected for further experimental characterization strains EA-12 and EABMAA-21 as representatives of the *Pantoea* ‘OTU 7’ from *E. atala*; strains PF2-63 and PF-27 as representatives of the *Serratia* ‘OTU 52’ from *P. floridana*; and the strains PFBMAA-4 and RS-48, as representative of the *Stenotrophomonas* ‘OTU 792’ and ‘OTU 6’ from *P. floridana* and *R. slossoni*, respectively. Interestingly, these latter strains represent two distinct lineages within the *Stenotrophomonas* genus. These OTUs, which we propose to be keystone species, are either present or ubiquitous in the microbiomes of all insect species from the global sampling (**Fig. 3B, Table S10**). We also identified and isolated strains of the previously mentioned uncultured *Klebsiella*-like genus (**Fig. 3A**), which we propose to also be a keystone taxa. Further analysis of these isolates was not pursued due to our inability to obtain representative MAGs from the sub-community co-cultures and the uncertainty about their taxonomic classification needed for phylogenomics analyses.

### Metagenomic mining of natural products reveals a conserved BGC core

Metagenomes from the six sub-community co-cultures were analyzed with antiSMASH and found to contain a total of 262 predicted BGCs, of which 85 were complete and non-redundant. These included 16 different types of BGCs, the most abundant of which were biosynthetic systems for aryl polyenes and non-ribosomal peptides (NRPs), with 20 and 19 BGCs, respectively (20% of the total). The remaining BGCs were bacteriocins (8), terpenes (11), NRPS-independent siderophores (5), hybrid PKS-NRPs (3), thiopeptides (3), resorcinols (2), butyrolactone (2), Hserlactone (1), type III PKS (1), lasso peptide (1), acyl amino acids (1), type II PKS (1), ectoine (1), and others (6) (**Table S11**). Of these, two core BGCs were present in all *EcoMining* metagenomes from the three insects: one turnerbactin-like BGC from the catechol-type siderophore category, and one carotenoid-like aryl polyene (**Fig. S2, Table S12**).

Further analysis of these two core BGCs using BiG-SCAPE revealed that the turnerbactin-like and aryl polyene BGCs formed five and four clans, respectively, mostly composed of scaffolds assigned to *Pantoea* “OTU 7” mainly from the *E. atala* metagenomes, and *Serratia “*OTU 52” and *Stenotrophomonas* “OTU 6” and “OTU 792” mainly from the *R. slossoni* metagenomes. Other siderophore BGCs of the hydroxamate and mixed ligands types were found in *Stenotrophomonas* and other bacterial genera, such as *Acinetobacter, Klebsiella*, and *Enterobacter* (**Fig. 4A**). Most of the clans of either type of BGCs were composed of BGCs from only one bacterial taxon, with the exception that the biggest clan of aryl polyenes, which contained BGCs from both *Pantoea* and *Serratia*.

**Fig. 4.**
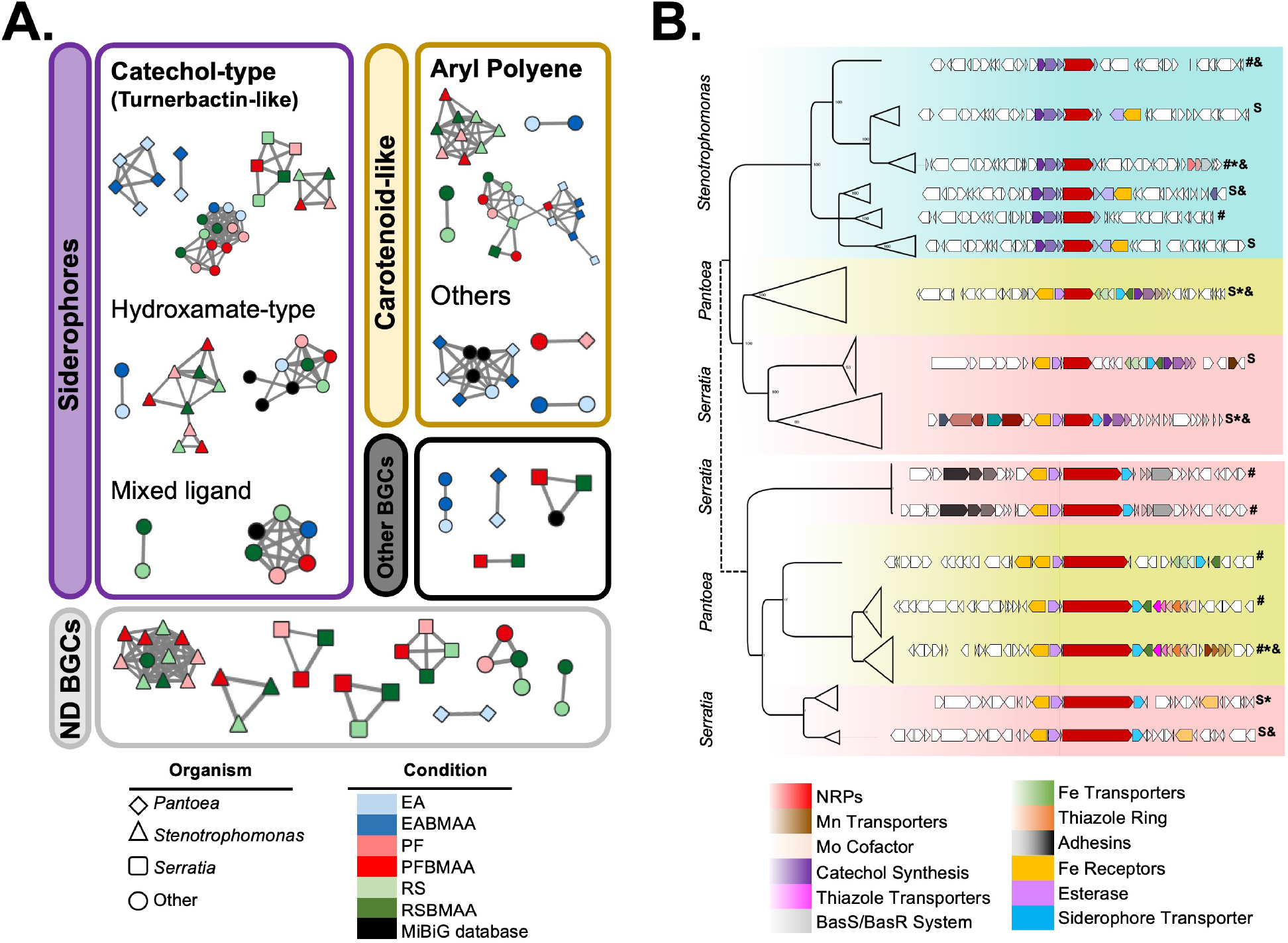
Conserved BGCs of cycadivorous core gut microbiome. **A**. BiG-SCAPE networks containing 85 complete and non-redundant BGCs identified in the six metagenomes obtained from the *EcoMining* experiments. Turnerbactin-like, catechol-type siderophore, and aryl polyene BGCs formed five and four clans, respectively. **B**. CORASON analysis with 230 turnerbactin-like BGCs from *Serratia, Pantoea*, and *Stenotrophomonas* genomes were used to reconstruct a BGC phylogeny. Turnerbactin-like BGCs can be separated into two groups based on changes in the NRPs’ protein sequences and their genomic context. The first group was found in all three bacterial genera, while the second was found only in *Serratia* and *Pantoea*. Two types of divergently related BGCs were found: (i) *bona fide* catechol-type siderophore BGCs (marked with S), and (ii) siderophore-like BGCs, no Fur box detected (#). BGCs found in the isolated strains from co-cultures are denoted with asterisk (*), and BGCs found in MAGs with ampersands (&). Expanded versions of the trees supporting this general phylogeny are provided as supplementary information (**Fig. S7, S8, S9, and S10**).

### Phylogenomics and semi-untargeted metabolomics of siderophores reveals specialized metabolism convergence

To explore the evolutionary dynamics of the biosynthetic potential of core and putative keystone taxa, whole BGC CORASON phylogenies and core proteome species phylogenies were obtained. The *Pantoea* phylogeny indicates that *Pantoea* sp. EA-12 and *Pantoea* sp. EABMAA-21 are part of a monophyletic clade that includes *Pantoea* species previously isolated from plants and rivers, while *Pantoea* sp. Pa-EAmG belongs to a monophyletic clade of *Pantoea* isolated from insects, soil, and plants (**Fig. S3**). In the *Serratia* phylogeny, *Serratia* sp. PF2-63 and *Serratia* sp. PF-27 formed a monophyletic clade with MAGs *Serratia* sp. Se-RSBMAAmG, *Serratia* sp. Se-PFmG, *Serratia* sp. Se-PFBMAAmG, and *Serratia* sp. Se-RSmG, and *Serratia* sp. OMLW3, an isolate from the insect, *Orius majusculus* (Chen et al., 2017) (**Fig. S4**). The *Stenotrophomonas* phylogeny was composed of two monophyletic clades, one containing the genomes and MAGs from *Stenotrophomonas* sp. RS-48, *Stenotrophomonas* sp. St-RSmG, and *Stenotrophomonas* sp. St-RSBMAAmG, plus isolates from soils and rivers; and the other composed of *Stenotrophomonas* sp. PFBMAA-4, *Stenotrophomonas* sp. St-PFmG, and *Stenotrophomonas* sp. St-PFBMAAmG (**Fig. S5**). These phylogenies place the strains proposed as keystone taxa within an evolutionary context specific to the insect’s gut, and provide a framework to look into BGCs distribution.

A contrasting evolutionary dynamic was found for the two conserved BGCs from the previous section. On one hand, the carotenoid-like aryl polyene BGC was highly conserved across the phylogenies of the core and putative keystone genera *Pantoea* “OTU 7”, *Serratia* “OTU 52”, and *Stenotrophomonas* “OTU 6 and OTU 792”, with changes limited to taxonomic distance (**Fig. S6**). In contrast, the turnerbactin-like BGCs, for which in-depth functional annotation of each locus was performed, were composed of two types of divergently related BGCs: (i) *bona fide* catechol-type siderophore BGCs and (ii) siderophore-like BGCs for which some of the expected key functional elements for a siderophore BGC, e.g., regulatory iron boxes could not be detected (**Fig. S7, S8, S9, and S10**). This analysis also showed that turnerbactin-like BGCs could be separated into two groups based on changes in the NRP protein sequences and their genomic vicinity. The first group was found in all three bacterial genera, while the second was found only in *Serratia* and *Pantoea* (**Fig. 4B**). The gene composition of the BGCs – i.e., the presence or absence of genes for recognition of certain metals such as Fe, Mn, and Mo – further separated the turnerbactin-like BGCs found in each genus.

To explore the chemical diversification suggested by these phylogenomics analyses, Global Natural Products Social (GNPS) molecular networks (Aron et al., 2020) were constructed using MS data obtained from pooled peaks collected after HPLC separation **(Fig. S11)**. To ensure that metal-chelating metabolites were targeted, we adopted a differential growth condition and analytical sample preparation with + / - ferric iron, as we have previously done in other microbial ecology studies (Cruz-Morales et al., 2017). Identification of metal chelating metabolites revealed 9 clusters of related compounds, with 7 of these metabolites conserved between the three bacterial core genera *Pantoea, Serratia*, and *Stenotrophomonas* (**Fig. 5A, Table S13**). Among these, we identified as conserved the siderophores ferrioxamine H ([M-2H+Fe]^+^= 514.1717 *m/z*, cluster 1, **Fig. 5B**) and B ([M-2H+Fe]^+^= 614.2715 *m/z*, cluster 3, **Fig. 5C**), which are the succinylated dimer and the acetyl-capped trimer, respectively. Unexpectedly, however, a putative desferrioxamine (*des*) BGC was not detected in any of the assembled metagenomes of the co-cultures, nor in the genomes that were sequenced independently. Our unsuccessful genome mining efforts included antiSMASH analyses; direct mapping of the reads of all six genome sequencing libraries against the *des* BGC from a close relative known to produce desferrioxamines, *Pantoea agglomerans* (MiBiG ID: BGC0001572); and identification of the Fur-dependent iron boxes implicated in iron metabolism regulation and previously exploited for siderophores genome mining (Fillat, 2014). The results of the latter analysis identified loci potentially involved in iron metabolism, of which some included biosynthetic enzymes and/or receptors, but were dissimilar to the DesABCD enzymes or the DesE receptor (**Table S14**). Together, these results provide evidence that metal-chelating metabolites are diversifying in the gut microbiome of cycad-feeding insects.

**Fig. 5.**
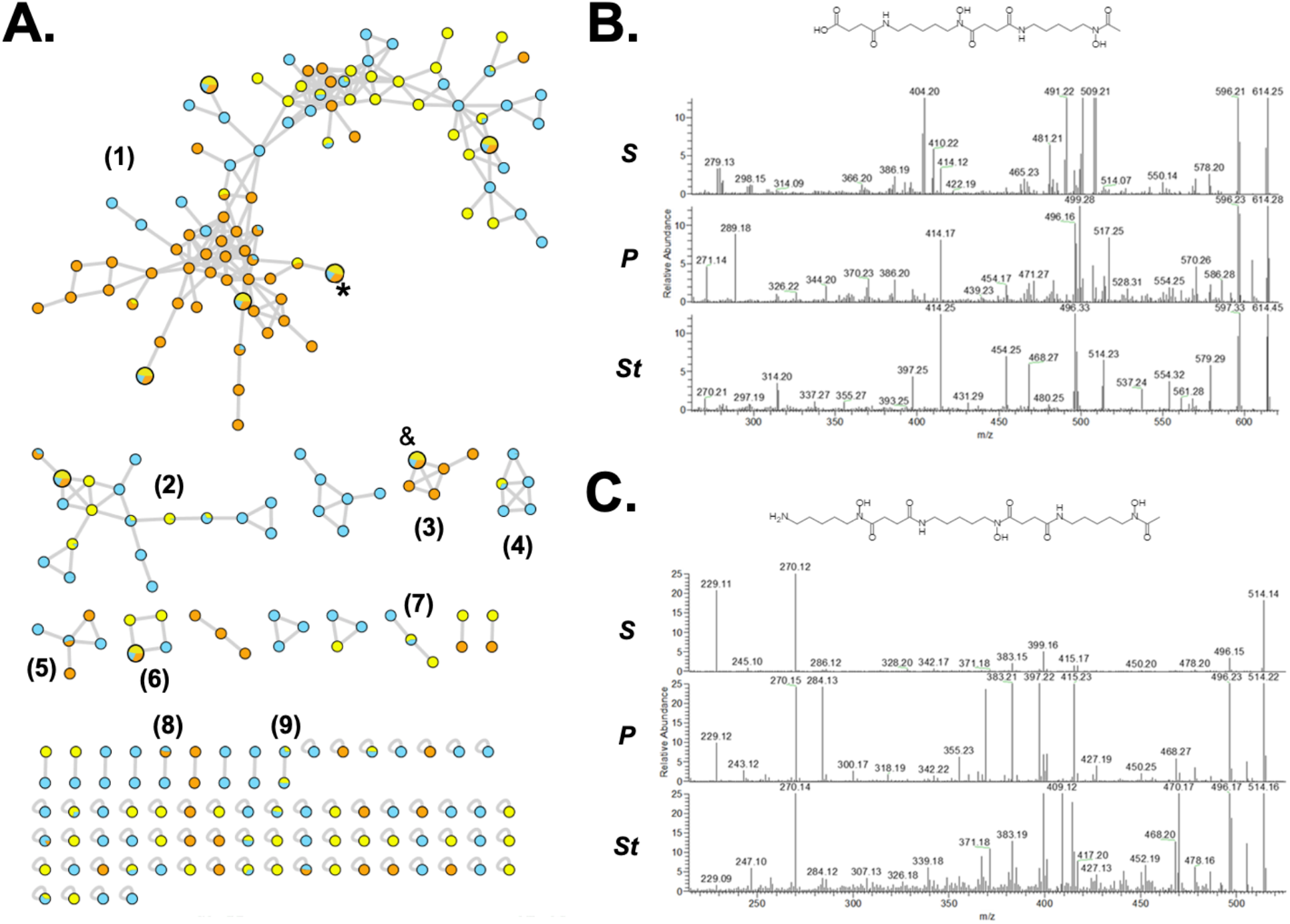
Conserved chemically divergent siderophores present in keystone taxa. **A**. GNPS networks revealed 9 conserved clusters of metal-chelating metabolites produced by keystone bacteria of the genera *Pantoea* (Blue), *Serratia* (Yellow), and *Stenotrophomonas* (Orange). Four of these clusters included at least one metabolite shared amongst all three genera, but cluster 1 had four of these metabolites. Ferrioxamines H (*) and B (&) were identified in cluster 1 and 3, respectively, after a combined manual and GNPS annotation. Selected metabolites of each group, and their MS features, are provided in **Table S13. B**. Ferrioxamine H and **C**. Ferrioxamine B. MS/MS spectra of these known siderophores was manually analyzed and compared with standards and databases to confirm the validity of the GNPS annotation. *S* = *Serratia, P* = *Pantoea*, and *St* = *Stenotrophomonas*.

## Discussion

According to the 16S community surveys, the core gut microbiomes of Coleoptera tended to be richer than those of Lepidoptera, which is consistent with previous reports that lepidopteran larvae typically do not host large resident bacterial communities (Whitaker et al., 2016; Hammer et al., 2017). However, we found that a cross-species core microbiome is widespread among cycadivorous insects – including several lepidopterans – regardless of geographic location, insect order, life stage, host use, or feeding guild. Several of these core bacteria belong to clades that engage in beneficial and pathogenic associations with plants and insects, such as *Pantoea* and *Stenotrophomonas*. Although it is possible that similar microbiome convergence may be a more widespread feature of insect faunas that feed on chemically distinctive host plants, most insect microbiome surveys focus on a single species or group rather than on the microbiota derived from a community of herbivores, making it difficult to compare our results to those from similarly specialized insect faunas.

Co-culturing of microbial sub-communities, the key feature of the *EcoMining* approach, revealed that BMAA does not affect bacterial community composition. This is to be expected, as the insects’ gut lumens should contain BMAA from the insects’ food plants, and any bacteria that cannot survive in BMAA-rich environments are likely naturally excluded from these communities. Genome and metagenome mining revealed a rich biosynthetic potential, including a set of core BGCs found in all three insect species. Although only three insect species were included in the BGC analysis, all 12 insect species from the global 16S survey contained the same bacterial OTUs from which the core BGCs were found, suggesting that the specific biosynthetic potential observed in the gut microbiomes of *Eumaeus, Rhopolotria*, and *Pharaxanotha* might be widespread among cycadivorous insects.

Further BGC analysis recovered and confirmed a high diversity of metal-chelating metabolites, e.g., siderophores, which are low molecular weight metabolites secreted by bacteria to chelate metals, often in response to iron deficiency, but which also play many ecologically relevant roles within bacterial communities (Kramer et al., 2020). Several catechol-type siderophores were predicted from the genomes of bacteria known to be enterobactin-like siderophore producers, such as *Pantoea* and *Stenotrophomonas* (Hisatomi et al., 2021; Reitz et al., 2017). Based on this evidence, the availability of iron (and potentially also other metals) may drive the diversification of these specialized metabolites within the insects’ gut microbiomes, as was recently shown in *Serratia plymuthica* with its combinatorial siderophore biosynthetic potential from polyamines (Cleto et al., 2021). Given that previous research into insect-siderophore interactions is limited, it is unclear whether the rich metal-chelating diversity observed within the microbiomes of cycadivorous is a widespread feature of insect gut microbiomes, or if it is limited to insects with specialized diets. Semi-untargeted metabolomics and molecular networking of metal-chelating metabolites produced by keystone taxa showed signs of chemical diversification, consistent with the genetic diversity revealed by genome and metagenome mining. Carotenoid-like aryl polyene biosynthetic systems, which were found to be conserved, remain to be investigated but it is interesting to note that they have been implicated in honeybee resistance to pathogenic fungi (Miller et al., 2021).

It is unknown how siderophores in the guts of cycadivorous insects might impact host fitness. A beneficial role of siderophores has been observed in plants, whereby siderophores produced by commensal or mutualistic bacteria can reduce growth of plant pathogens (Kupferschmied, Maurhofer, & Keel, 2013). While research into siderophore-insect interactions is limited, previous studies have detected bacterial siderophores in the guts of mosquitoes, honey bees, grasshoppers, and moths (Ganley et al., 2020; Hertlein, 2014; Indiragandhi, 2008; Sonawane, 2018). In each of these cases, however, siderophore production was implicated in bacterial pathogenicity with clear costs for host fitness. Indeed, for many phyto- and entomopathogenic bacteria, siderophore production contributes to host colonization and microbial virulence, for example by competing with the host for iron and other essential micronutrients that are typically limited in plants. Without detailed assessments of the availability of iron and other metals to cycadivorous insects, it is unclear whether siderophore-producing bacteria may similarly compete with their hosts for iron.

An interesting possibility is that siderophore-producing bacteria could provide protective benefits for cycadivorous insects by mediating chemical interactions between metal ions and BMAA inside the gut lumen. BMAA is also capable of chelating iron and other metals (Glover et al., 2012), and the formation of BMAA-metal ion complexes has been proposed as one possible cause of neurotoxicity in humans (Diaz-Parga et al., 2018, 2021; Nunn et al., 1989). The chelation of divalent metal ions alters the equilibrium between BMAA and its corresponding carbamate adducts, which are required for binding to glutamate receptors and exerting neurotoxic effects (Diaz-Parga et al., 2021). Within the insect gut, the affinity of siderophores, such as desferrioxamines, for metal ions is likely to be higher than that of BMAA, such that bacterial siderophores may compete with BMAA for limited iron (and/or other metal ions serving as enzyme co-factors) within the insect gut lumen. In turn, this would alter the equilibrium between BMAA and its carbamate adducts affecting BMAA’s toxicity within the gut. This hypothesis warrants further investigation.

Chemical diversification of bacterial specialized metabolites in the guts of cycadivorous insects is likely the result of complex ecological and evolutionary forces, including the biomolecular and physicochemical features of natural products synthesized by the insect and/or plant hosts, and may lead to novel and unexpected metabolic functions (Chevrette et al., 2020). Within this context, iron-limitation is likely not the only factor driving siderophore production within the insects’ bacterial communities. Instead, we hypothesize that metal-chelating metabolites, e.g., siderophores, serve other functions besides chelating metals as a means for nutrient acquisition, as recently suggested in other systems (Shepherdson & Elliot, 2022). That this genetic and chemical diversity has been selected for in bacterial taxa found across all 12 of the insects we surveyed suggests that these specialized metabolites are a key feature of cycadivorous insects’ gut microbiomes, and perhaps of their feeding ecology. Future studies that take a plant-centered approach to surveying insect microbiomes, in combination with functional investigations such as the *EcoMining* approach implemented here, may ultimately yield more insight into the role of insects’ gut microbiomes in facilitating convergent adaptation to specific hosts.

## Acknowledgments

We are grateful to Jose Luis Steffani-Vallejo, Juan D. Camarena-Alba, and Paulina Mejía-Ponce for their help in laboratory work, and Nelly Selem-Mojica, Diego Garfias-Gallegos, María Fernanda Contreras-González, Hannah Augustijn for bioinformatics support. Thanks also to Hermann Staude and Horace Tan for their invaluable contributions to fieldwork, to William Tang and Irene Terry for providing additional insect specimens, and Rory Maher for making data collection possible. This study was supported by CONACyT #169701 and FON.INST./265/2016 to A.C.J. CONACyT #179290, #177568 and #285746 to F.B.G., also supported by a Newton Advanced Fellowship of The Royal Society, UK (NAF/R2/180631). National Science Foundation (NSF) of the US #1541560 to N.E.P. M.R.L.W was supported by NSF of the US Postdoctoral Research Fellowship in Biology (1309425), with fieldwork funded by the Harvard Museum of Comparative Zoology’s Putnam Expeditionary Fund, the Wildlife Reserves Singapore Conservation Fund, and the Explorer’s Club. S.S. was supported by NSF of the US Graduate Research Fellowship and Postdoctoral Research Fellowship in Biology (1906333).

