## Supplementary_Figures for "Specialized metabolic convergence in the gut microbiomes of cycad-feeding insects tolerant to β-methylamino-L-alanine (BMAA)"

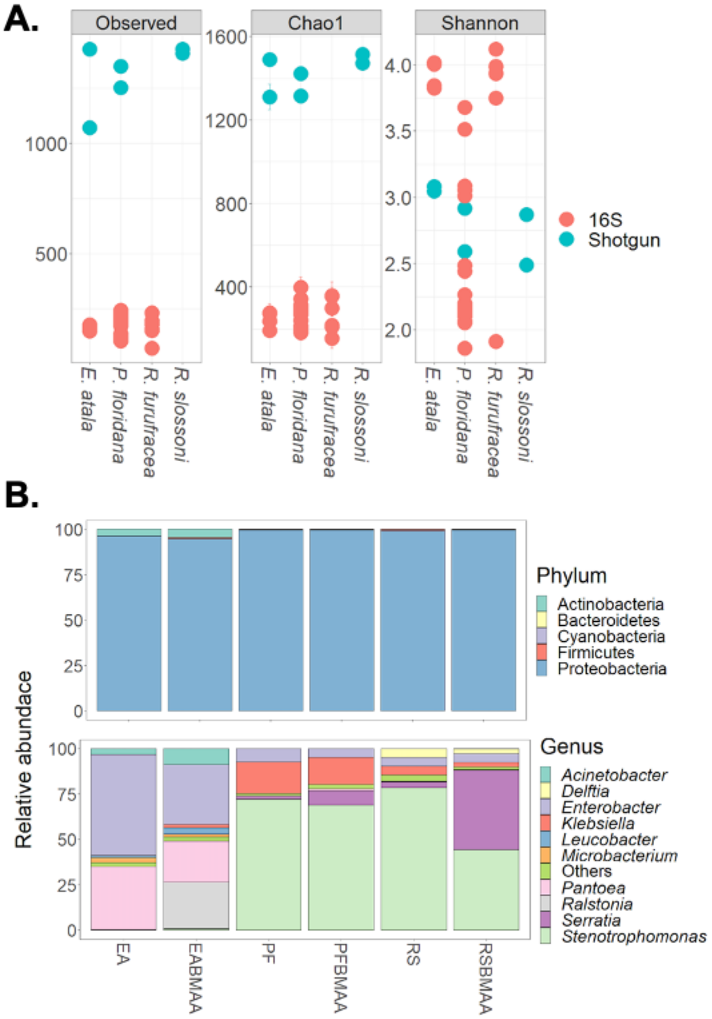


**Supplementary Figure S1. Taxonomic analysis of shotgun metagenomes. A.** Alpha diversity comparison of 16S and shotgun metagenomes. **B.** Relative abundance of taxonomically classified and filtered reads in each shotgun metagenome assigned to the phylum and genus level, and the 10 most abundant genera.

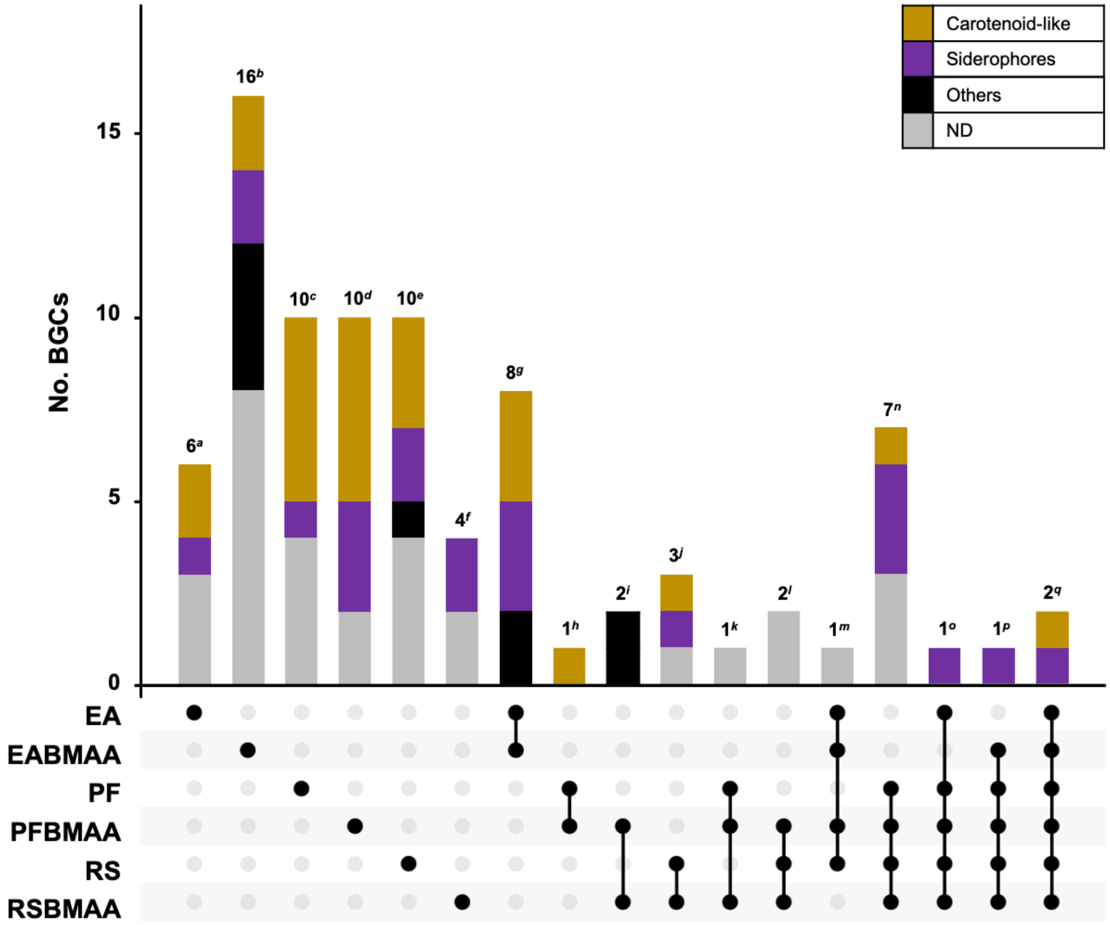


**Supplementary Figure S2. Presence and absence BGC plot.** 85 complete and non-redundant BGCs identified in the six metagenomes obtained from the co-cultures were used to construct the BGC plot. Three specific BGCs, one turnebactin-like BGC from the catechol-type siderophore category and two carotenoid-like aryl polyenes, were found to be present in all the metagenomes.

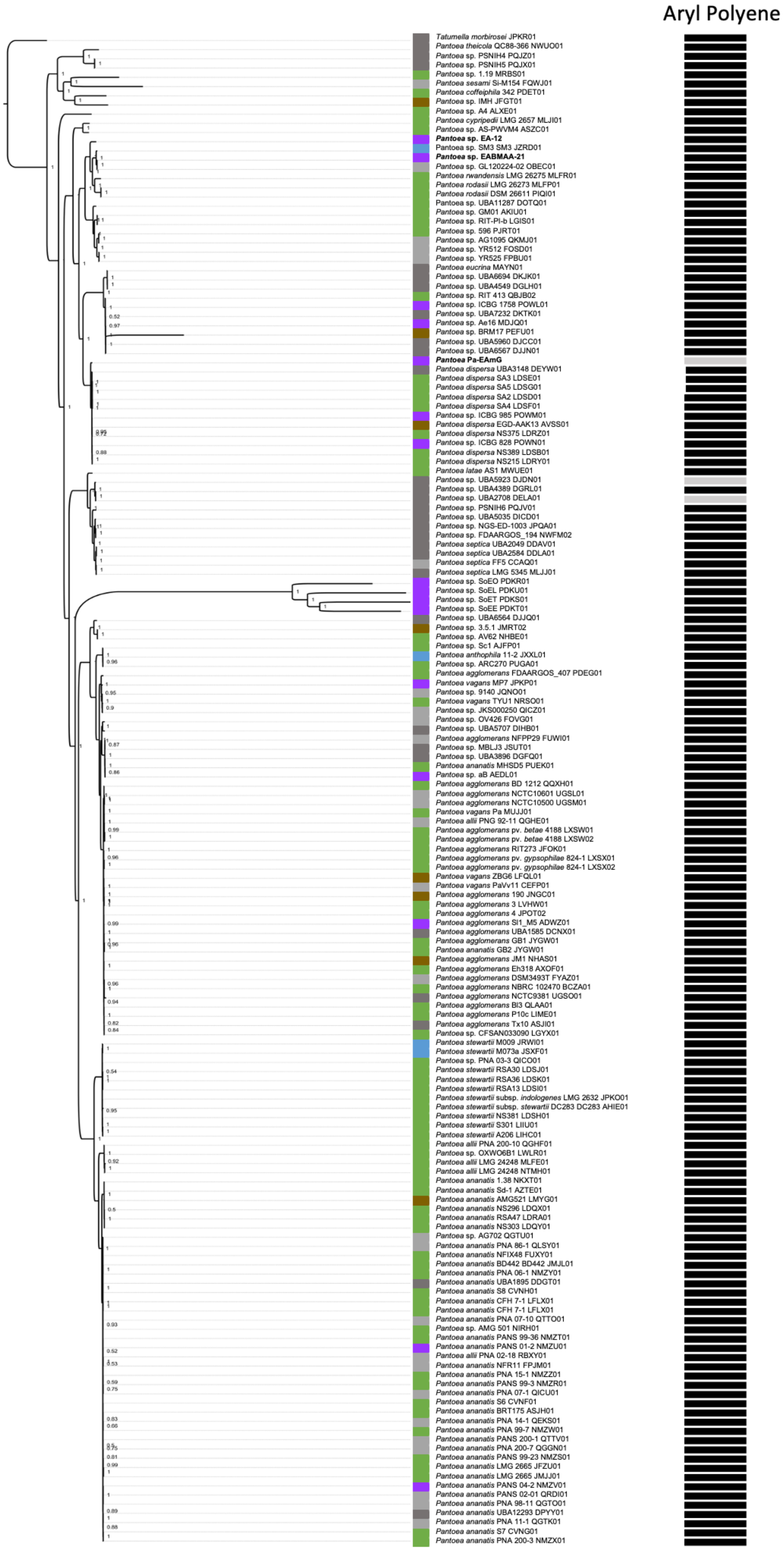
Aryl Polyene
Tatumella morbirosei JPKR01
Pantoea theicola QC88-366 NWUO01
Pantoea sp. PSNIH4 PQJZ01
Pantoea sp. PSNIH5 PQJX01
Pantoea sp. 1.19 MRBS01
Pantoea sesami Si-M154 FQWJ01
Pantoea coffeiphila 342 PDET01
Pantoea sp. IMH JFGT01
Pantoea sp. A4 ALXE01
Pantoea cypripedii LMG 2657 MLJI01
Pantoea sp. AS-PWVM4 ASZC01
Pantoea sp. EA-12
Pantoea sp. SM3 SM3 JZRD01
Pantoea sp. EABMAA-21
Pantoea sp. GL120224-02 OBEC01
Pantoea rwandensis LMG 26275 MLFR01
Pantoea rodasii LMG 26273 MLFP01
Pantoea rodasii DSM 26611 PIQI01
Pantoea sp. UBA11287 DOTQ01
Pantoea sp. GM01 AKIU01
Pantoea sp. RIT-PI-b LGIS01
Pantoea sp. 596 PJRT01
Pantoea sp. AG1095 QKMJ01
Pantoea sp. YR512 FOSD01
Pantoea sp. YR525 FPBU01
Pantoea eucrina MAYN01
Pantoea sp. UBA6694 DKJK01
Pantoea sp. UBA4549 DGLH01
Pantoea sp. RIT 413 QBJB02
Pantoea sp. ICBG 1758 POWL01
Pantoea sp. UBA7232 DKTK01
Pantoea sp. Ae16 MDJQ01
Pantoea sp. BRM17 PEFU01
Pantoea sp. UBA5960 DJCC01
Pantoea sp. UBA6567 DJJN01
Pantoea Pa-EAmG
Pantoea dispersa UBA3148 DEYW01
Pantoea dispersa SA3 LDSE01
Pantoea dispersa SA5 LDSG01
Pantoea dispersa SA2 LDSD01
Pantoea dispersa SA4 LDSF01
Pantoea sp. ICBG 985 POWM01
Pantoea dispersa EGD-AAK13 AVSS01
Pantoea dispersa NS375 LDRZ01
Pantoea sp. ICBG 828 POWN01
Pantoea dispersa NS389 LDSB01
Pantoea dispersa NS215 LDRY01
Pantoea latae AS1 MWUE01
Pantoea sp. UBA5923 DJDN01
Pantoea sp. UBA4389 DGRL01
Pantoea sp. UBA2708 DELA01
Pantoea sp. PSNIH6 PQJV01
Pantoea sp. UBA5035 DICD01
Pantoea sp. NGS-ED-1003 JPQA01
Pantoea sp. FDAARGOS_194 NWFM02
Pantoea septica UBA2049 DDAV01
Pantoea septica UBA2584 DDLA01
Pantoea septica FF5 CCAQ01
Pantoea septica LMG 5345 MLJJ01
Pantoea sp. SoEO PDKR01
Pantoea sp. SoEL PDKU01
Pantoea sp. SoET PDKS01
Pantoea sp. SoEE PDKT01
Pantoea sp. UBA6564 DJJQ01
Pantoea sp. 3.5.1 JMRT02
Pantoea sp. AV62 NHBE01
Pantoea sp. Sc1 AJFP01
Pantoea anthophila 11-2 JXXL01
Pantoea sp. ARC270 PUGA01
Pantoea agglomerans FDAARGOS_407 PDEG01
Pantoea vagans MP7 JPKP01
Pantoea sp. 9140 JQNO01
Pantoea vagans TYU1 NRSO01
Pantoea sp. JKS000250 QICZ01
Pantoea sp. OV426 FOVG01
Pantoea sp. UBA5707 DIHB01
Pantoea agglomerans NFPP29 FUWI01
Pantoea sp. MBLJ3 JSUT01
Pantoea sp. UBA3896 DGFQ01
Pantoea ananatis MHSD5 PUEK01
Pantoea sp. aB AEDL01
Pantoea agglomerans BD 1212 QQXH01
Pantoea agglomerans NCTC10601 UGSL01
Pantoea agglomerans NCTC10500 UGSM01
Pantoea vagans Pa MUJJ01
Pantoea allii PNG 92-11 QGHE01
Pantoea agglomerans pv. betae 4188 LXSW01
Pantoea agglomerans pv. betae 4188 LXSW02
Pantoea agglomerans RIT273 JFOK01
Pantoea agglomerans pv. gypsophilae 824-1 LXSX01
Pantoea agglomerans pv. gypsophilae 824-1 LXSX02
Pantoea vagans ZBG6 LFQL01
Pantoea vagans PaVv11 CEFP01
Pantoea agglomerans 190 JNGC01
Pantoea agglomerans 3 LVHW01
Pantoea agglomerans 4 JPOT02
Pantoea agglomerans SI1_M5 ADWZ01
Pantoea agglomerans UBA1585 DCNX01
Pantoea agglomerans GB1 JYGW01
Pantoea ananatis GB2 JYGW01
Pantoea agglomerans JM1 NHAS01
Pantoea agglomerans Eh318 AXOF01
Pantoea agglomerans DSM3493T FYAZ01
Pantoea agglomerans NBRC 102470 BCZA01
Pantoea agglomerans NCTC9381 UGSO01
Pantoea agglomerans BI3 QLAA01
Pantoea agglomerans P10c LIME01
Pantoea agglomerans Tx10 ASJI01
Pantoea sp. CFSAN033090 LGYX01
Pantoea stewartii M009 JRWI01
Pantoea stewartii M073a JSXF01
Pantoea sp. PNA 03-3 QICO01
Pantoea stewartii RSA30 LDSJ01
Pantoea stewartii RSA36 LDSK01
Pantoea stewartii RSA13 LDSI01
Pantoea stewartii subsp. indologenes LMG 2632 JPKO01
Pantoea stewartii subsp. stewartii DC283 DC283 AHIE01
Pantoea stewartii NS381 LDSH01
Pantoea stewartii S301 LIIU01
Pantoea stewartii A206 LIHC01
Pantoea allii PNA 200-10 QGHF01
Pantoea sp. OXWO6B1 LWLR01
Pantoea allii LMG 24248 MLFE01
Pantoea allii LMG 24248 NTMH01
Pantoea ananatis 1.38 NKXT01
Pantoea ananatis Sd-1 AZTE01
Pantoea ananatis AMG521 LMYG01
Pantoea ananatis NS296 LDQX01
Pantoea ananatis RSA47 LDRA01
Pantoea ananatis NS303 LDQY01
Pantoea sp. AG702 QGTU01
Pantoea ananatis PNA 86-1 QLSY01
Pantoea ananatis NFIX48 FUXY01
Pantoea ananatis BD442 BD442 JMJL01
Pantoea ananatis PNA 06-1 NMZY01
Pantoea ananatis UBA1895 DDGT01
Pantoea ananatis S8 CVNH01
Pantoea ananatis CFH 7-1 LFLX01
Pantoea ananatis CFH 7-1 LFLX01
Pantoea ananatis PNA 07-10 QTTO01
Pantoea sp. AMG 501 NIRH01
Pantoea ananatis PANS 99-36 NMZT01
Pantoea ananatis PANS 01-2 NMZU01
Pantoea allii PNA 02-18 RBXY01
Pantoea ananatis NFR11 FPJM01
Pantoea ananatis PNA 15-1 NMZZ01
Pantoea ananatis PANS 99-3 NMZR01
Pantoea ananatis PNA 07-1 QICU01
Pantoea ananatis S6 CVNF01
Pantoea ananatis BRT175 ASJH01
Pantoea ananatis PNA 14-1 QEKS01
Pantoea ananatis PNA 99-7 NMZW01
Pantoea ananatis PANS 200-1 QTTV01
Pantoea ananatis PNA 200-7 QGGN01
Pantoea ananatis PANS 99-23 NMZS01
Pantoea ananatis LMG 2665 JFZU01
Pantoea ananatis LMG 2665 JMJJ01
Pantoea ananatis PANS 04-2 NMZV01
Pantoea ananatis PANS 02-01 QRDI01
Pantoea ananatis PNA 98-11 QGTO01
Pantoea ananatis UBA12293 DPYY01
Pantoea ananatis PNA 11-1 QGTK01
Pantoea ananatis S7 CVNG01
Pantoea ananatis PNA 200-3 NMZX01

**Supplementary Figure S3. Full *Pantoea* phylogenetic tree of representative strains, MAGs and isolated strains from the co-cultures.** 168 *Pantoea* genomes were used to reconstruct this phylogeny using the core proteome composed of 64 proteins (Table S5 and S8). Habitats for each species are indicated with colored bullets. Purple = insects, Green = plants, Brown = soil, Blue = water, Dark gray = Other, and Light gray = Not determined. The incidence of the aryl polyene BGCs is shown as presence (black bars) or absence (light gray bars).

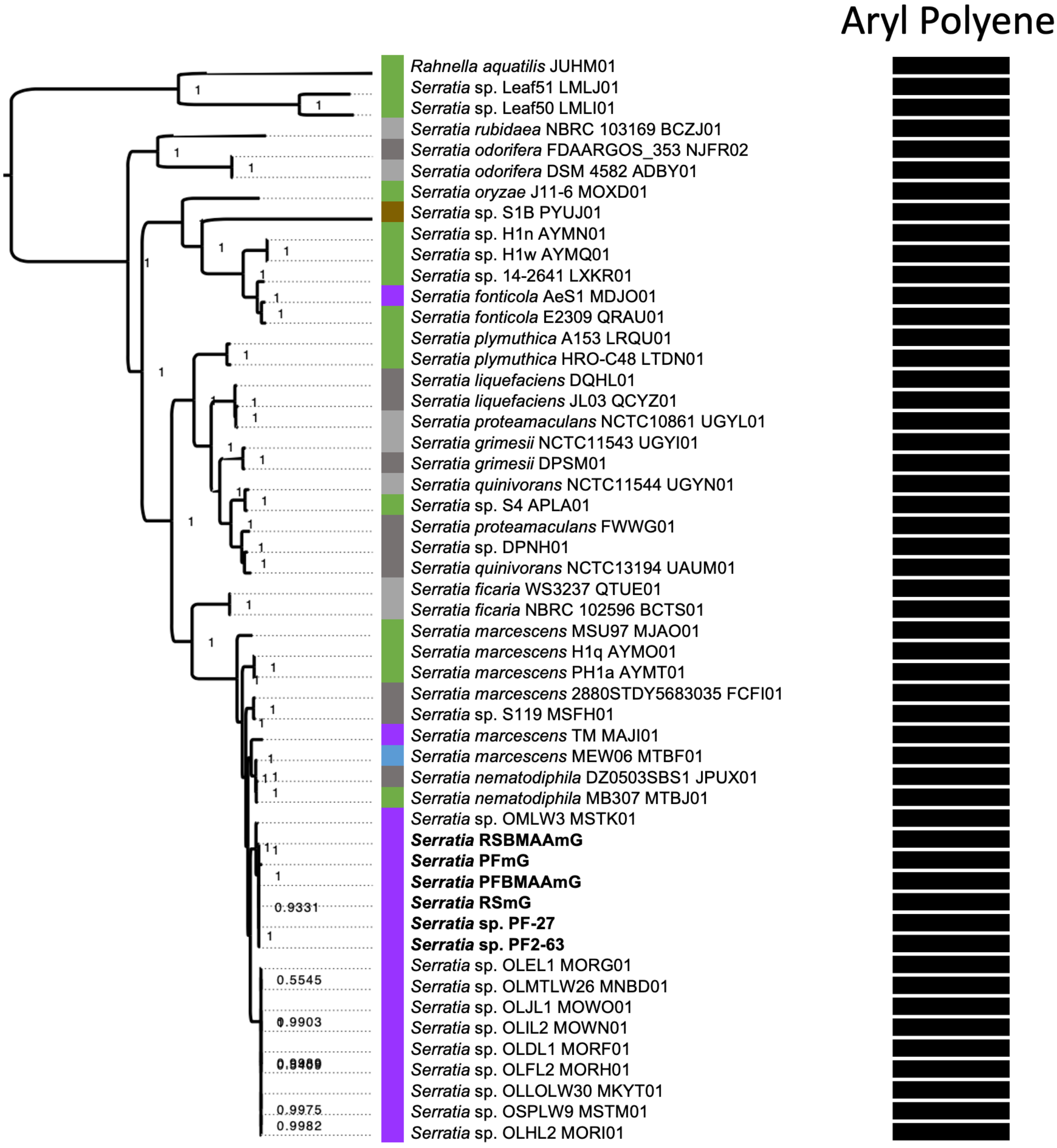


**Supplementary Figure S4. Full *Serratia* phylogenetic tree of representative strains, MAGs and isolated strains from the co-cultures.** 51 *Serratia* genomes were used to reconstruct this phylogeny using the core proteome composed of 712 proteins (Table S6 and S9). Habitats for each species are indicated with colored bullets. Purple = insects, Green = plants, Brown = soil, Blue = water, Dark gray = Other, and Light gray = Not determined. The incidence of the aryl polyene BGCs is shown as present in all the *Serratia* genomes (Black bars).

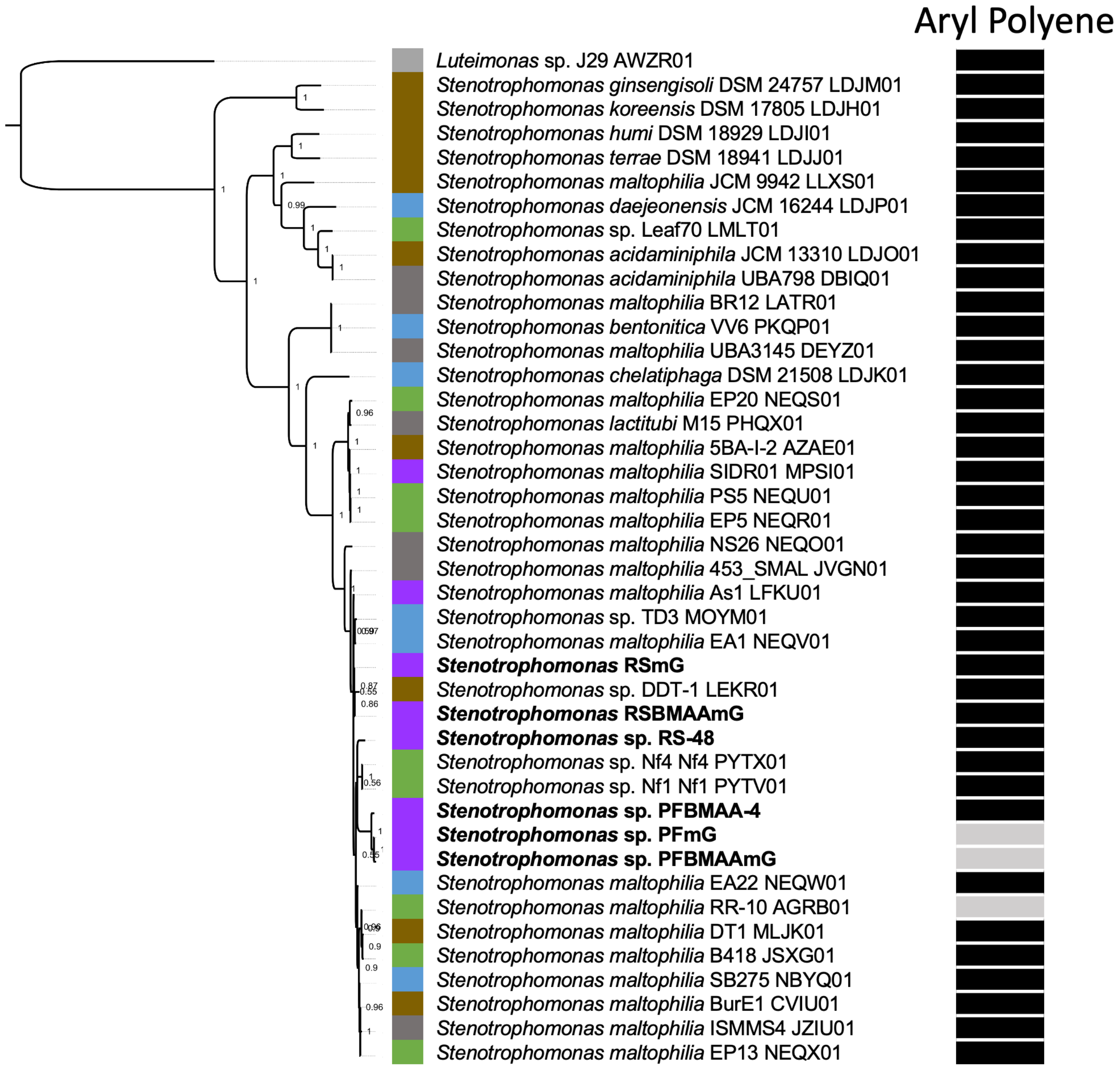


**Supplementary Figure S5. Full *Stenotrophomonas* phylogenetic tree of representative strains, MAGs and isolated strains from the co-cultures.** 41 *Serratia* genomes were used to reconstruct this phylogeny using the core proteome composed of 39 proteins (Table S4 and S7). Habitats for each species are indicated with colored bullets. Purple = insects, Green = plants, Brown = soil, Blue = water, Dark gray = Other, and Light gray = Not determined. The incidence of the aryl polyene BGCs is shown as presence (black bars) or absence (light gray bars).

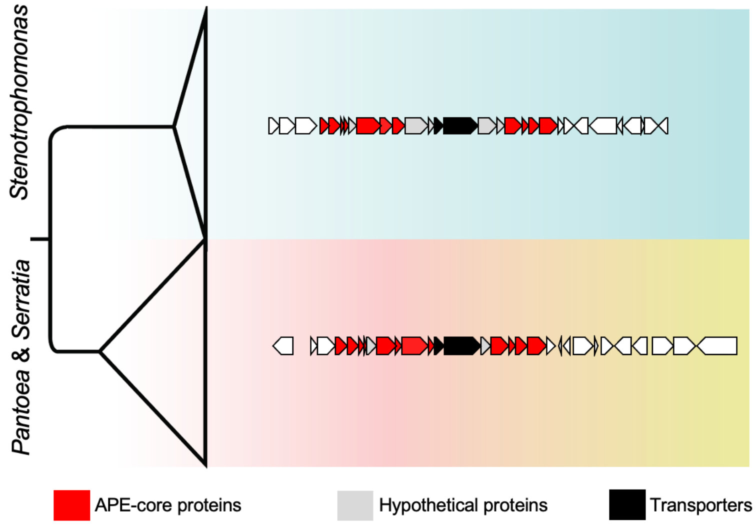


**Supplementary Figure S6. Aryl Polyene BGCs phylogeny of *Serratia, Pantoea,* and *Stenotrophomonas* species.** 254 Aryl polyene BGCs from S*erratia, Pantoea*, and *Stenotrophomonas* genomes were used to reconstruct this phylogeny. Aryl polyene BGCs are highly conserved in all three bacterial genera.

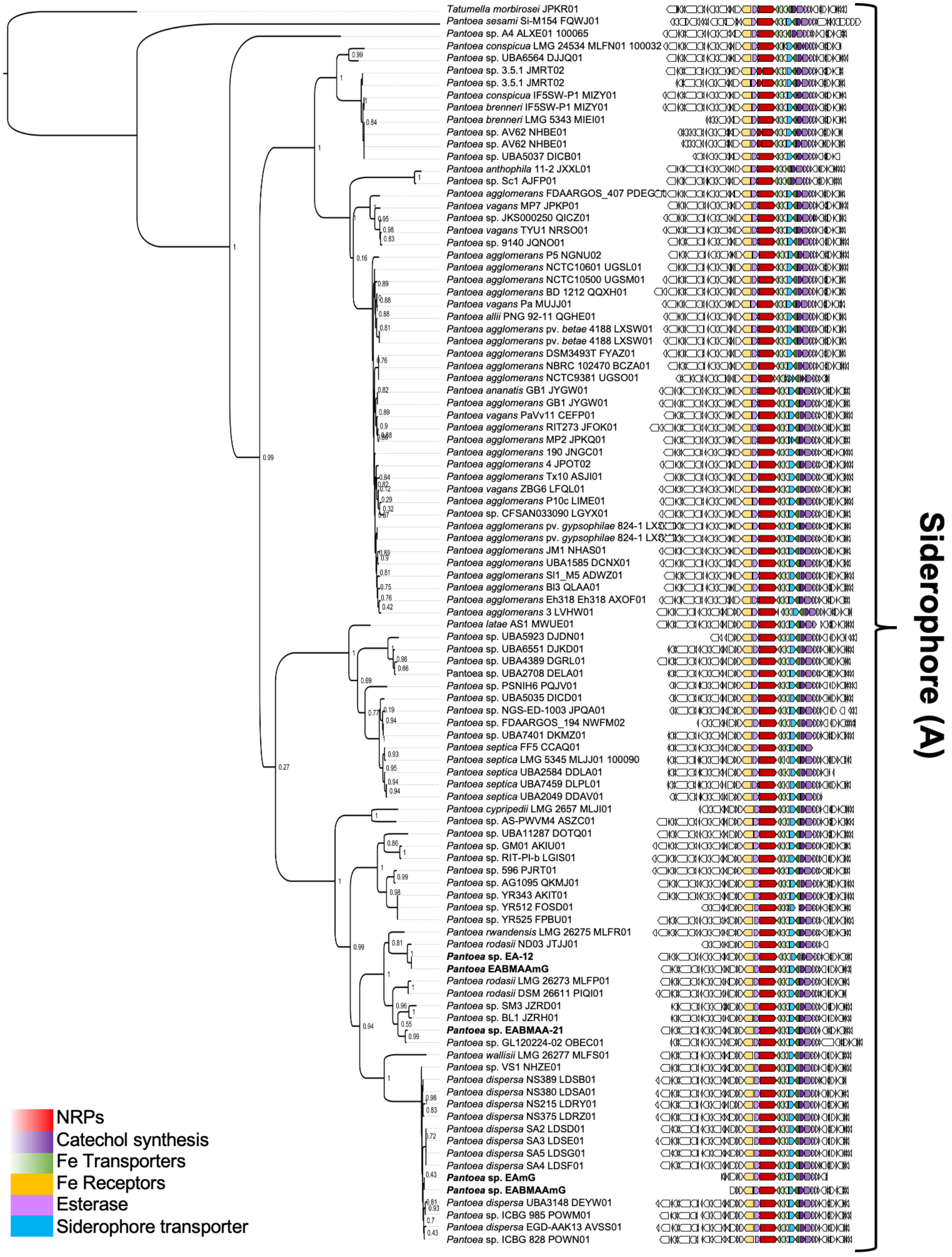
Tatumella morbirosei JPKR01
Pantoea sesami Si-M154 FQWJ01
Pantoea sp. A4 ALXE01 100065
Pantoea conspicua LMG 24534 MLFN01 100032
Pantoea sp. UBA6564 DJJQ01
Pantoea sp. 3.5.1 JMRT02
Pantoea sp. 3.5.1 JMRT02
Pantoea conspicua IF5SW-P1 MIZY01
Pantoea brenneri IF5SW-P1 MIZY01
Pantoea brenneri LMG 5343 MIEI01
Pantoea sp. AV62 NHBE01
Pantoea sp. AV62 NHBE01
Pantoea sp. UBA5037 DICB01
Pantoea anthophila 11-2 JXXL01
Pantoea sp. Sc1 AJFP01
Pantoea agglomerans FDAARGOS_407 PDEG
Pantoea vagans MP7 JPKP01
Pantoea sp. JKS000250 QICZ01
Pantoea vagans TYU1 NRSO01
Pantoea sp. 9140 JQNO01
Pantoea agglomerans P5 NGNU02
Pantoea agglomerans NCTC10601 UGSL01
Pantoea agglomerans NCTC10500 UGSM01
Pantoea agglomerans BD 1212 QQXH01
Pantoea vagans Pa MUJJ01
Pantoea allii PNG 92-11 QGHE01
Pantoea agglomerans pv. betae 4188 LXSW01
Pantoea agglomerans pv. betae 4188 LXSW01
Pantoea agglomerans DSM3493T FYAZ01
Pantoea agglomerans NBRC 102470 BCZA01
Pantoea agglomerans NCTC9381 UGSO01
Pantoea ananatis GB1 JYGW01
Pantoea agglomerans GB1 JYGW01
Pantoea vagans PaVv11 CEFP01
Pantoea agglomerans RIT273 JFOK01
Pantoea agglomerans MP2 JPKQ01
Pantoea agglomerans 190 JNGC01
Pantoea agglomerans 4 JPOT02
Pantoea agglomerans Tx10 ASJI01
Pantoea vagans ZBG6 LFQL01
Pantoea agglomerans P10c LIME01
Pantoea sp. CFSAN033090 LGYX01
Pantoea agglomerans pv. gypsophilae 824-1 LX
Pantoea agglomerans pv. gypsophilae 824-1 LX
Pantoea agglomerans JM1 NHAS01
Pantoea agglomerans UBA1585 DCNX01
Pantoea agglomerans SI1_M5 ADWZ01
Pantoea agglomerans BI3 QLAA01
Pantoea agglomerans Eh318 Eh318 AXOF01
Pantoea agglomerans 3 LVHW01
Pantoea latae AS1 MWUE01
Pantoea sp. UBA5923 DJDN01
Pantoea sp. UBA6551 DJKD01
Pantoea sp. UBA4389 DGRL01
Pantoea sp. UBA2708 DELA01
Pantoea sp. PSNIH6 PQJV01
Pantoea sp. UBA5035 DICD01
Pantoea sp. NGS-ED-1003 JPQA01
Pantoea sp. FDAARGOS_194 NWFM02
Pantoea sp. UBA7401 DKMZ01
Pantoea septica FF5 CCAQ01
Pantoea septica LMG 5345 MLJJ01 100090
Pantoea septica UBA2584 DDLA01
Pantoea septica UBA7459 DLPL01
Pantoea septica UBA2049 DDAV01
Pantoea cypripedii LMG 2657 MLJI01
Pantoea sp. AS-PWVM4 ASZC01
Pantoea sp. UBA11287 DOTQ01
Pantoea sp. GM01 AKIU01
Pantoea sp. RIT-PI-b LGIS01
Pantoea sp. 596 PJRT01
Pantoea sp. AG1095 QKMJ01
Pantoea sp. YR343 AKIT01
Pantoea sp. YR512 FOSD01
Pantoea sp. YR525 FPBU01
Pantoea rwandensis LMG 26275 MLFR01
Pantoea rodasii ND03 JTJJ01
Pantoea sp. EA-12
Pantoea EABMAAmG
Pantoea rodasii LMG 26273 MLFP01
Pantoea rodasii DSM 26611 PIQI01
Pantoea sp. SM3 JZRD01
Pantoea sp. BL1 JZRH01
Pantoea sp. EABMAA-21
Pantoea sp. GL120224-02 OBEC01
Pantoea wallisii LMG 26277 MLFS01
Pantoea sp. VS1 NHZE01
Pantoea dispersa NS389 LDSB01
Pantoea dispersa NS380 LDSA01
Pantoea dispersa NS215 LDRY01
Pantoea dispersa NS375 LDRZ01
Pantoea dispersa SA2 LDSD01
Pantoea dispersa SA3 LDSE01
Pantoea dispersa SA5 LDSG01
Pantoea dispersa SA4 LDSF01
Pantoea sp. EAmG
Pantoea sp. EABMAAmG
Pantoea dispersa UBA3148 DEYW01
Pantoea sp. ICBG 985 POWM01
Pantoea dispersa EGD-AAK13 AVSS01
Pantoea sp. ICBG 828 POWN01
NRPs
Catechol synthesis
Fe Transporters
Fe Receptors
Esterase
Siderophore transporter
Siderophore (A)

**Supplementary Figure S7. Turnerbactin-like BGC phylogeny of *Pantoea*.** 101 turnerbactin-like BGCs were used to reconstruct this phylogeny using the conserved proteins present in all BGCs. Genomic context visualization, as well as in-deep functional annotation of each BGC, reveal highly conservation in all the organisms.

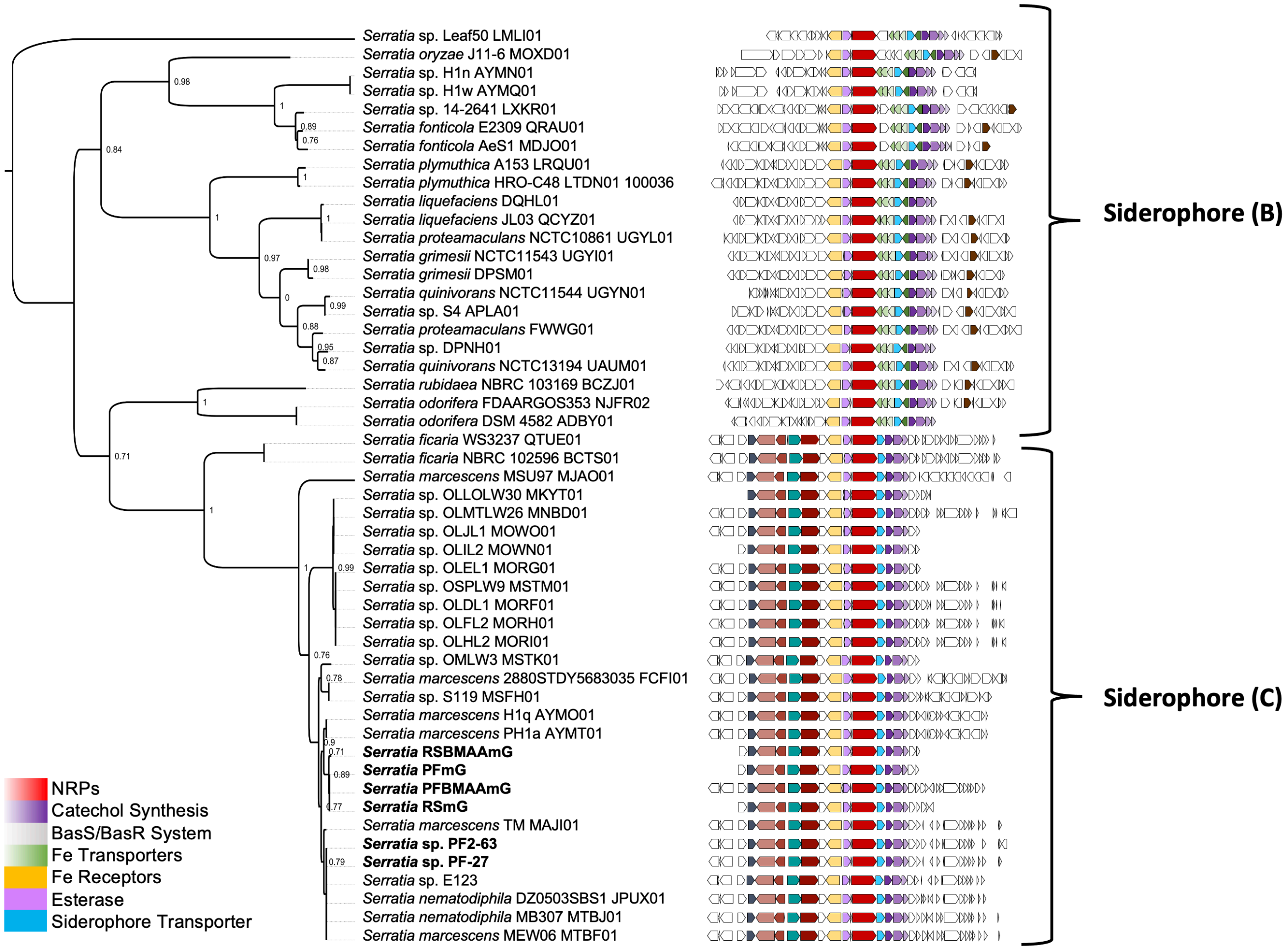


**Supplementary Figure S8. Turnerbactin-like BGC phylogeny of *Serratia*.** 50 turnerbactin-like BGCs were used to reconstruct this phylogeny using the conserved proteins present in all BGCs. Genomic context visualization, as well as in-deep functional annotation of each BGC, reveal two different catechol-type BGCs. Siderophore (C) BGC is present in all the *Serratia* (meta)genomes obtained from cycadivorous guts.

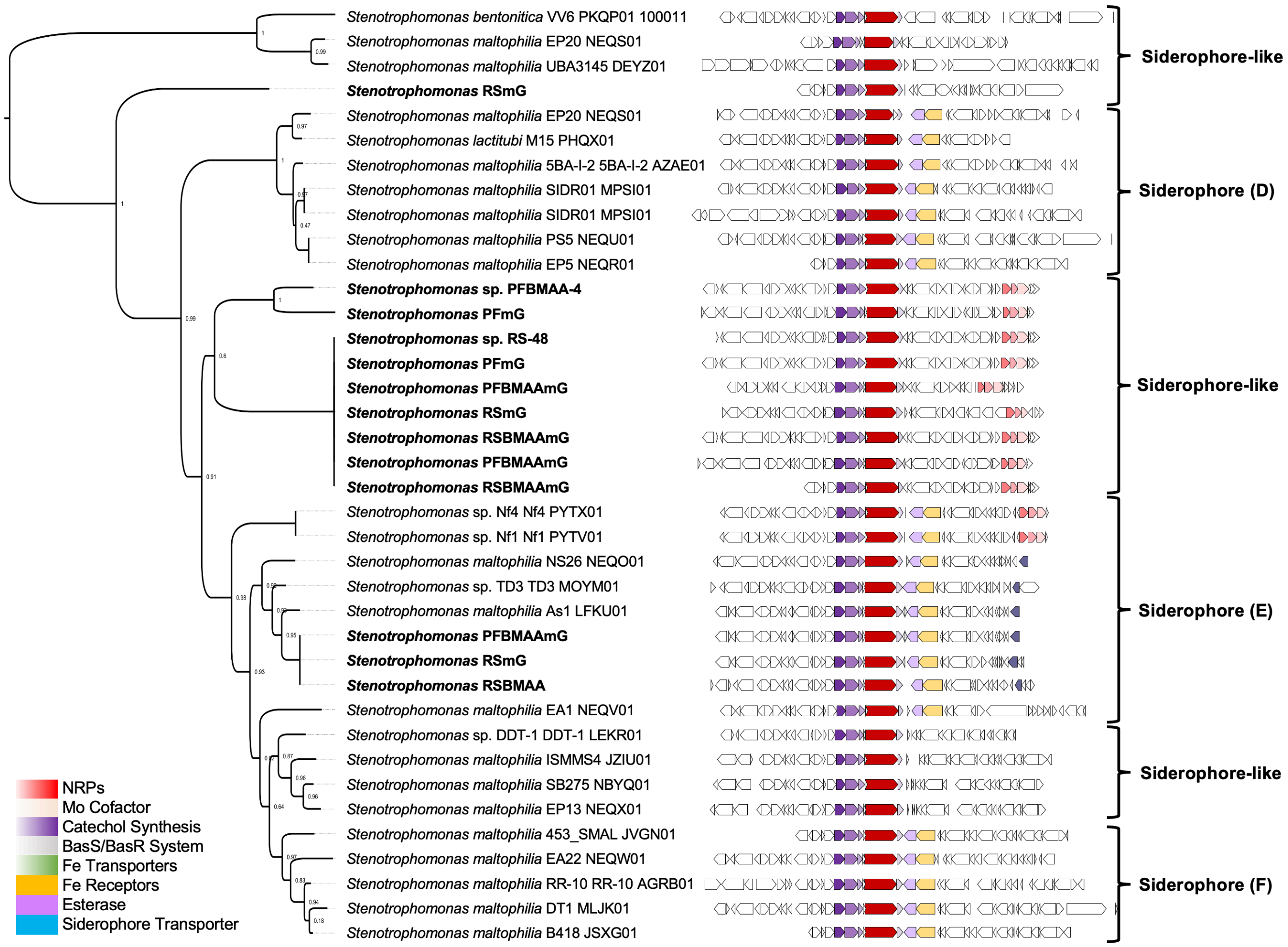


**Supplementary Figure S9. Turnerbactin-like BGC phylogeny of *Stenotrophomonas*.** 38 turnerbactin-like BGCs were used to reconstruct this phylogeny using the conserved proteins present in all BGCs. Genomic context visualization, as well as in-deep functional annotation of each BGC, reveal three *bona fide* catechol-type BGCs (D, E, and F), plus three siderophore-like BGCs, some of them present in *Stenotrophomonas* (meta)genomes obtained from cycadivorous guts.

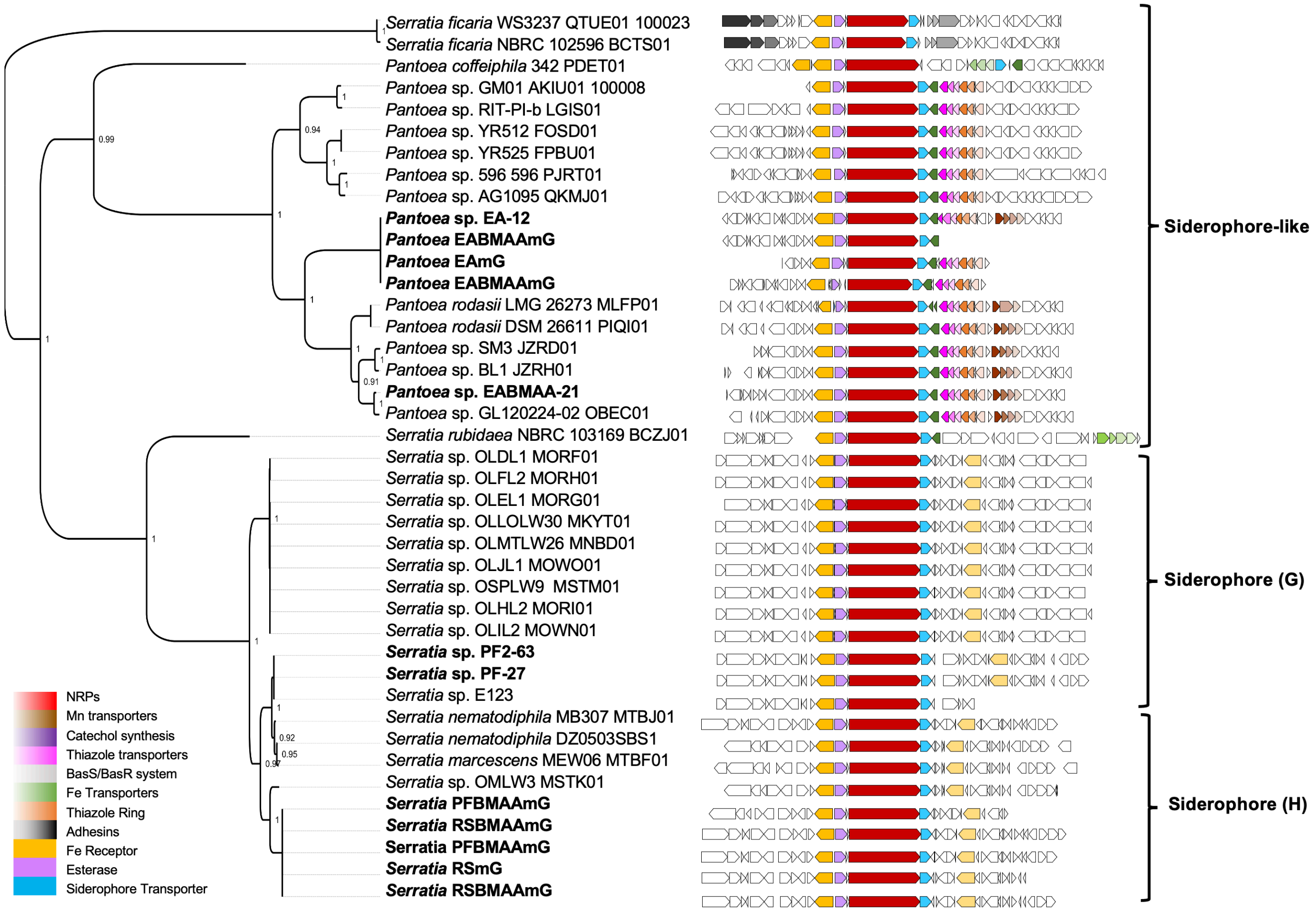


**Supplementary Figure S10. Turnerbactin-like BGC phylogeny of *Serratia-Pantoea*.** 41 enterobactin-like BGCs from both *Serratia* and *Pantoea* genomes were used to reconstruct this phylogeny. Genomic context visualization, as well as in-deep functional annotation of each BGC, reveal one siderophore-like BGC present in some *Serratia* and *Pantoea* genomes plus a catechol-type BGC (G) present exclusively in *Serratia* (meta)genomes.

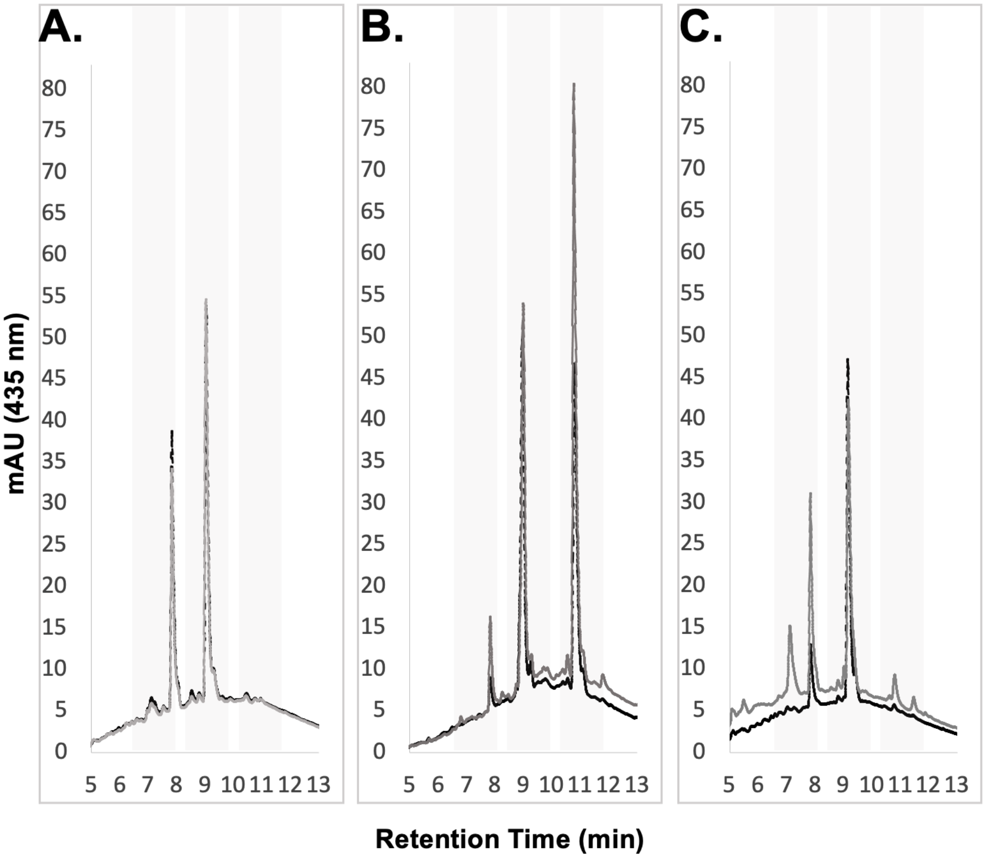


**Supplementary Figure S11. Identification of siderophores produced by six bacterial strains isolated from cycadivorus insects through HPLC.** HPLC analysis of **A.** Two *Serratia* strains: PF2-63 (dash line) and PF-27 (solid line), **B.** Two *Pantoea* strains: EA-12 (dash line) and EABMAA-21 (solid line), and **C.** Two *Stenotrophomonas:* PFBMAA-4 (dash line) and RS-48 (solid line) under siderophore-promoting conditions revealed signals at 435 nm associated with the production of these compounds. The indicated retention times (gray selection) were then collected and analyzed by MS-MS mass spectrometry.
